# Schistosome esophageal gland factor MEG-8.2 drives host cell lysis and interacts with host immune proteins

**DOI:** 10.1101/2024.11.15.623777

**Authors:** Pallavi Yadav, Sabona B. Simbassa, Ryan Sloan, Phillip A. Newmark, Jayhun Lee

## Abstract

Schistosomes are blood flukes that ingest large amounts of host blood during their intra-mammalian stage. The ingested blood contains leukocytes that can be harmful, yet the parasites survive inside the host for decades, reflecting superb immune evasion mechanisms that remain poorly understood. Our previous work discovered that FoxA, a forkhead transcription factor, drives the production of the esophageal gland, an anterior digestive organ essential for degrading the ingested leukocytes and for *in vivo* survival. However, a comprehensive molecular makeup of the esophageal gland remains unclear. Importantly, which of the esophageal gland factors are responsible for degrading the ingested leukocytes, their mechanism of action, and how such a function relates to parasite survival and immune evasion remains unknown. Here, we identify additional esophageal gland genes by taking a comparative transcriptomics approach to identify transcripts altered in *foxA* knockdown adult schistosomes. A targeted RNAi screen coupled with biochemistry reveals that specific domains of the micro-exon gene MEG-8.2, can drive host cell lysis in a concentration-dependent manner. Using pull-down assays coupled with mass spectrometry, we discover that MEG-8.2 interacts with several host membrane and extracellular proteins that play important roles in activating innate and/or adaptive immunity. Together, our findings suggest a dual role for MEG-8.2 in effectively lysing the ingested cells in the esophageal lumen and interacting with specific host proteins to neutralize or suppress the host immunity. These findings lay an important foundation for exploiting esophageal gland factors to treat schistosomiasis.

## Introduction

During intramammalian homeostasis, adult schistosomes reproduce inside the host vasculature, producing hundreds of eggs daily, causing schistosomiasis (*1*). Schistosomiasis remains one of the most prevalent parasitic diseases, with over 200 million individuals affected globally (*2*). However, other than praziquantel, no treatment or preventative options are available, highlighting the urgent need to devise alternative approaches to target these parasites. Contrary to schistosome eggs, which are highly immunogenic, worms inside the host bloodstream can withstand attacks by the immune system, leading to their longevity. Previous studies have revealed the importance of the tegument, a syncytial outer skin of the parasite, to play a crucial role in this process. The tegument is structurally unique: the parasite-host interface is composed of a double lipid bilayer, a feature uniquely found among the blood flukes (*3*). In addition, host glycolipids and glycoproteins are found on the tegument, suggesting that their acquisition might help the parasite avoid detection by the host immune system (*4*). More recently, parasite stem cells have become the prime suspect in the successful development, homeostasis, reproduction, and propagation across the life cycle (*5, 6*). In adults, a significant proportion of somatic stem cells differentiate into tegument cells via transcription factors such as Zfp-1-1, p53-1, and Klf4 to replenish the tegument, which undergoes high cellular turnover (*7–10*). Similar mechanisms have been reported for tissues underlying parasites’ digestive tract, in which transcription factor Hnf4 (hepatocyte nuclear factor 4) plays a crucial role in gut cell production and maintenance (*11*). Hnf4 knockdown parasites display increased *eled*+ (stem/ progenitor) cells with a decreased output of *ctsb*+ gut cells that perturbs digestion of red blood cells, resulting in parasite death *in vivo*.

Given that schistosomes consume large amounts of host blood containing potentially harmful immune components, these parasites likely deploy immune-evasion mechanisms through their digestive tract. Anterior to the gut is the parasites’ esophagus, which is surrounded by a digestive organ called the esophageal gland (EG). The EG has been suggested as the initial site of blood processing where ingested erythrocytes are broken down and damaged leukocytes appear tethered to the esophageal lumen (*12*). There are ∼1000 cells in an adult male EG (*12*). These cells are densely packed, and their cytoplasm extends into the esophageal lumen for secretion. Electron microscopy shows secretory granules and vesicles found throughout the cytoplasm. These cytoplasmic extensions form a 2-dimensional plate-like structure that runs longitudinally from anterior to posterior of the lumen (*12*). These plates are regularly spaced, creating a large luminal surface area for processing ingested blood. These observations suggest that the cell types and genes that comprise the EG orchestrate secretion and maintain tissue integrity and function.

Our recent work discovered an essential regulator of EG cell development and maintenance, a forkhead transcription factor, FoxA (*13*). Expression of *foxA* is enriched in the EG and its neighboring *h2b*+ stem/progenitor cells. Knockdown in adult schistosomes disrupted the expression of several EG genes, preventing the parasites from blocking and degrading leukocytes in the esophagus. Such a function appears essential for evading the host immune system: parasites lacking the EG are rapidly cleared from the bloodstream of immunocompetent mice while they can survive inside immunocompromised mice. Previous studies identified several genes enriched in the EG. For instance, several EG genes were discovered by comparing the transcriptomes obtained from anterior and posterior halves of adult parasites and validating the expression of anterior-enriched genes using whole-mount mRNA in situ hybridization (WISH) (*14*). More recently, single-cell RNA-sequencing (scRNA-seq) of parasites from different life cycle stages captured a few dozen EG cells with a handful of genes (*11*). However, many of these have not been validated as *bona fide* EG genes. Importantly, their functional role in blocking or degrading the ingested blood cells and their role in parasite survival and parasite-host interaction remains to be determined. In this study, we take a comparative transcriptomics approach to comprehensively identify EG genes. We use RNAi to systematically screen for candidate EG factors that play a role in degrading the ingested leukocytes. Furthermore, we use a biochemical approach to dissect the functional domains of the top candidate factors that disrupt the host cell membrane and interact with a specific set of host proteins. Our findings lead us to propose a concentration-dependent dual role of the identified EG factor in degrading the host cells and interacting with host cell proteins with known functions in immune-mediated defense against pathogens. Together, this study establishes the foundation for deciphering the mechanism of action of EG-released factors and raises the possibility of targeting these factors as therapeutic avenues for treating schistosomiasis.

## Results

### *foxA* RNAi RNA-seq comprehensively identifies esophageal gland genes

We previously reported that *foxA* (Smp_331700) is enriched in the EG and colocalizes with a known EG gene, *meg-4*, and neighboring *h2b*+ stem/progenitor cells (*13*). Adult scRNA-seq revealed that *foxA* is enriched in *eled*+ neoblasts and *prom2*+ cells (*11*), which supports the notion that it acts upstream to regulate stem cell-to-EG cell differentiation. Indeed, *foxA* knockdown results in the apparent loss of the EG tissue, which was evidenced by the loss of *meg-4* expression via fluorescence in situ hybridization (FISH), loss of PNA lectin signal that is highly enriched in the EG, and downregulation of several known EG genes by qPCR (*13*). Therefore, we hypothesized that capitalizing on *foxA* knockdown worms would reveal genes expressed in the EG tissue. We collected RNA from three batches of adult male (n=∼20 per replicate) and female (n=∼30-40 per replicate) worms after ∼2 weeks of *foxA* knockdown *in vitro* (**Figure 1A**). The integrity of total RNA was verified, and the knockdown of several known EG genes was confirmed by qPCR (**Figure S1**). Differential expression analysis (fold change ≤ −2; false discovery rate ≤ 0.05) revealed significantly downregulated genes in *foxA* knockdown: 57 in males and 45 in females, 37 of which were shared. *foxA* was included in this list, confirming the consistent and significant knockdown across all batches of males and females (**Figure 1E and Table S1**). Based on their annotation, 36 commonly downregulated genes could be divided into six categories: microexon genes (MEGs), enzymes, secreted proteins/toxins, membrane proteins, a protease inhibitor, and hypothetical proteins.

**Figure 1.**
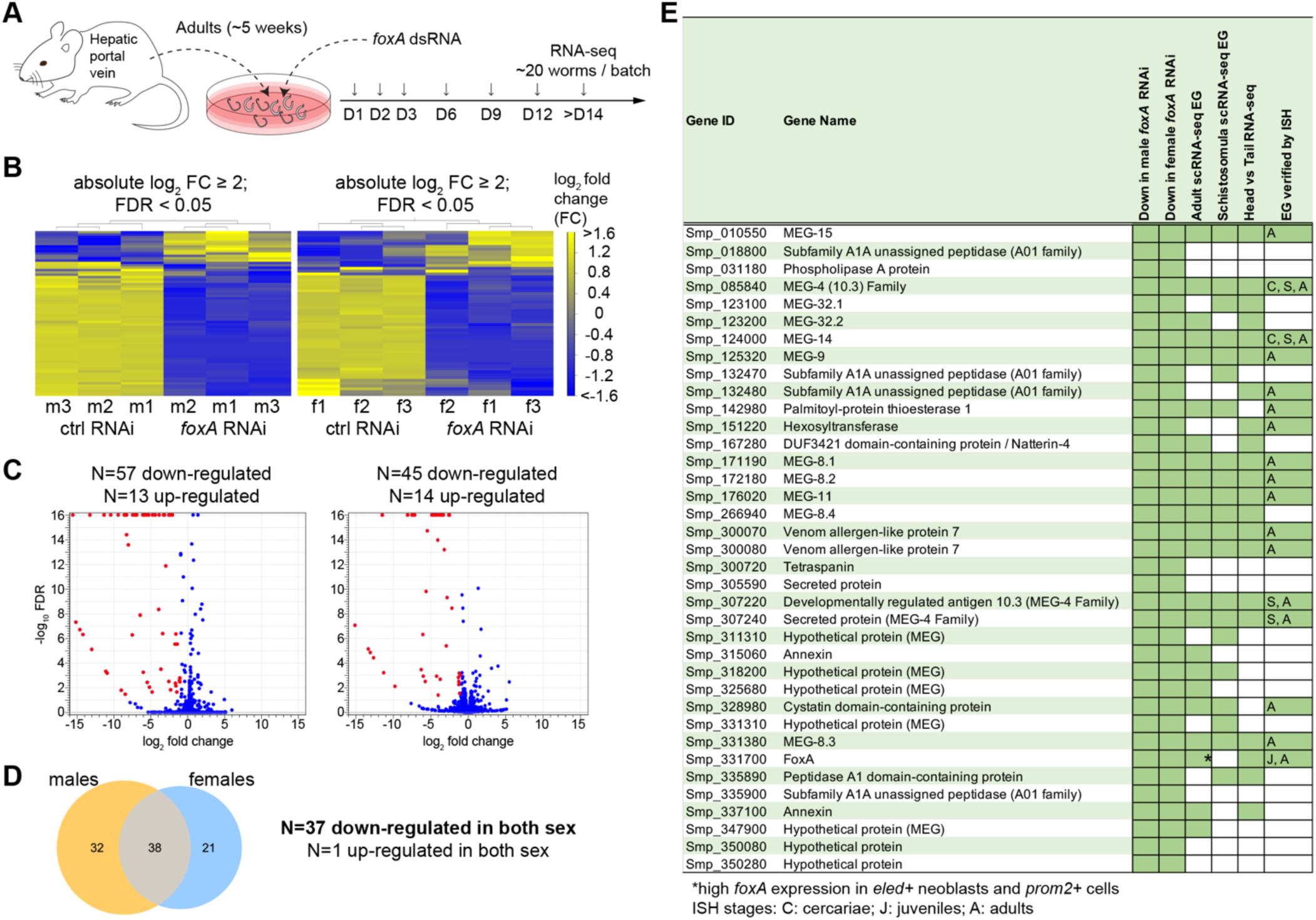
Comparative RNA-seq identifies genes downregulated in *foxA* knockdown parasites. (A) Experimental scheme to collect adult *foxA* RNAi parasites for RNA-sequencing. (B) Heatmap showing differentially expressed genes in males (left) and females (right). (C) Volcano plot of differentially expressed genes in males (left) and females (right). Red dots highlight downregulated genes. (D) Venn diagram shows 38 genes commonly differentially expressed in both sexes (37 downregulated and one upregulated). (E) Table of all commonly downregulated genes and their reported EG expression from adult scRNA-seq (*11*), schistosomula scRNA-seq (*19*), and head vs tail bulk RNA-seq (*14*). Those that are confirmed in their EG expression via *in situ* hybridization (ISH) are indicated with the analyzed stages.

50% (18 of 36) of the genes were microexon genes (MEGs). The protein-coding portion of the gene comprises microexons that typically range between 6 to 36 base pairs (*15, 16*). MEGs that are found in *Schistosomatidae* appear to have no similarity to other microexons in vertebrates (*17*) or aphids (*18*). Those previously validated by *in situ* hybridization as *bona fide* EG genes (*meg-4* family, *meg-8* family, *meg-9*, *meg-11*, *meg-14*, and *meg-15*) were among the top hits. The other MEGs in the list have also been reported to be expressed in the EG from either bulk (*14*) or schistosomula/adult scRNA-seq (*11, 19*), but their expression had not yet been validated. 22% (8 of 36) of the genes were enzymes such as *peptidase*, phospholipase A (*pla*), palmitoyl-protein thioesterase 1 (*ppt1*), and *hexosyltransferase*, most of which have been confirmed in their EG expression (*14, 16, 20*). In addition, known secreted proteins/toxins such as venom allergen-like protein 7 (*val-7*) and *natterin-4*, as well as a protease inhibitor *cystatin,* were identified in our dataset. These data suggest that our approach successfully identified the most commonly shared EG genes. Interestingly, several genes had never been reported to be in the EG, including *tsp* (Smp_300720), *secreted protein* (Smp_305590), *peptidase* (Smp_335900), and two *hypothetical proteins* (Smp_350080 and Smp_350280). We used colorimetric whole-mount *in situ* hybridization (WISH) to validate the EG expression of as many genes as possible (**Figure 2A**). In addition to known EG genes, we confirmed the enriched EG expression of *meg-32.1*, *meg-32.2*, *meg-8.4*, three other *meg*s (Smp_318200, Smp_325680, and Smp_347900), *natterin-4*, *annexin* (Smp_315060), and two peptidases (Smp_018800 and Smp_335890). *tsp* (Smp_300720) had a barely detectable signal in the EG, suggesting that RNA-seq captures even non-abundant transcripts. Outside of 37 shared downregulated genes, 20 were significantly downregulated in males and 8 were downregulated in females (**Figures 1C and 1D**). A few of these genes were enriched in the EG, such as *secreted protein* (Smp_331590) and *tsp* (Smp_320440) in both males and females (**Figure S2A**). Meanwhile, based on the published scRNA-seq atlas (*11*), other downregulated genes in males and females were enriched in non-EG cell types. For instance, several (4 of 8) genes downregulated in females included eggshell proteins that appear enriched in vitellocytes or eggs (**Figure S2B**). Similarly, several genes upregulated in *foxA* RNAi males (5 of 13) and females (2 of 14) were also related to reproductive development with vitellocytes and Mehlis gland enrichment. We speculate that these are likely due to the worms’ variable size and reproductive status and sparse ectopic Mehlis gland-forming males represented in different batches.

**Figure 2.**
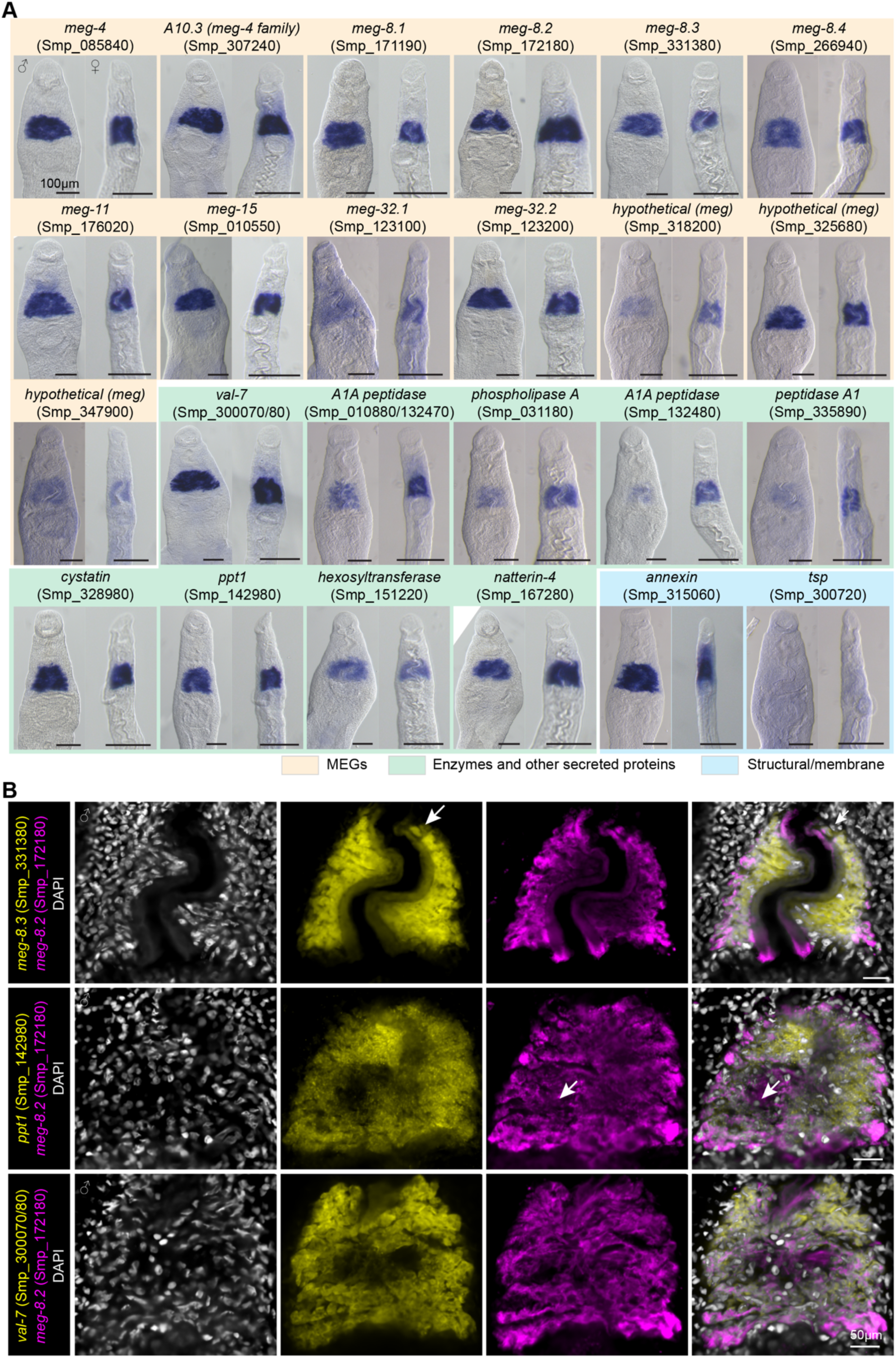
Genes downregulated in *foxA* knockdown are highly enriched in the EG. (A) WISH of downregulated genes in adult males and females. 5 to 10 worms per sex were analyzed for each gene. dFISH of select EG genes reveals subtle heterogeneous expression patterns. A few cells with a noticeable difference in signal between the two channels are indicated by the white arrow. A single confocal z-section from the head region of a male is shown for each combination. Between 5 to 10 males were analyzed per condition.

Having identified a core set of EG genes, we sought to determine if these genes are directly regulated by FoxA. Due to the lack of a specific antibody against FoxA, we took a bioinformatics approach by analyzing the local enrichment of binding motifs from the known transcription factor (JASPAR) database on the promoter sequences (5 kb upstream) of 36 genes (excluding *foxA*) using the MEME suite (*21*) (see Methods) (**Figures S3A – S3C**). Interestingly, among the 2430 motifs in the database, only nine motifs, all forkhead transcription factors, showed strong enrichment close to the transcription start site (∼600bp upstream) on 78% (28 of 36) of the promoter sequences. Moreover, the top two enriched motifs were those of human FOXE1 and FOXB1, which are homologous to Smp_342790 (*foxL*) and Smp_331700 (*foxA*). These results suggest a potential role of FoxA in directly regulating the transcription of core EG genes and that the expression of EG genes is likely associated with the differentiation of *foxA*+ stem cells.

To examine the potential heterogeneity of the EG cells, we used double fluorescence *in situ* hybridization (dFISH) of several identified markers (**Figure 2B**). Specifically, using *meg-8.2* (Smp_172180), we tested co-expression with another MEG (*meg-8.3*, Smp_331380) and two enzymes, *ppt1* (Smp_142980) and *val-7* (Smp_300070/80). We noticed a slight variation in signal intensity in some EG cells and a few cells that appeared to have noticeably low levels of one of the transcripts (white arrows), suggesting that heterogeneity may exist among the EG cells. However, virtually all EG cells expressed the analyzed EG markers, indicating that the impact of potential heterogeneity in the EG function might be low. Taken together, by exploiting *foxA* knockdown, we identified several new EG genes and confirmed multiple reported EG genes in their tissue-specific expression. Most identified genes have a conserved forkhead-binding motif in their promoter, suggesting that their expression is likely regulated directly by FoxA.

### FoxA is primarily restricted to the EG cell lineage, and EG loss does not perturb stem/progenitor cell balance

Adult scRNA-seq data suggests that *foxA* expression is highest in *eled*+ neoblasts and *prom2*+ intestinal progenitors (*11*). Previous work indicates that schistosome stem cells respond to tissue injury or cell death, increasing proliferation in efforts to replenish the missing cell types (*22*). *cbp1*, a CBP/p300 family transcriptional co-activator expressed throughout multiple stem/progenitor cell types, plays a role in this process throughout the body, including the EG. Thus, we sought to better understand the role of FoxA in stem cell-driven homeostasis. We analyzed our dataset to examine the expression changes of various stem/progenitor cell markers identified thus far from various single-cell studies (*11, 19, 23–27*) in *foxA* knockdown (**Figure S3D**). Expression levels of stem/progenitor cells’ subcluster markers, including *ago2-1*, *klf*, *nanos-2*, *fgfrA*, *fgfrB*, *hesl*, *zfp-1*, and *eled,* did not significantly change. Similarly, markers of *eled*-related subclusters (i.e., *astf* and *bhlh*), germline stem cells (*nanos-1*), S1 vitellocytes (*nr/vf1*), and somatic lineage progenitors (tegument: *p53-1* and *zfp-1-1*; gut: *hnf4*; flame cells: *sialidase*) also were not significantly different in *foxA* knockdown worms. *vwa*, a marker of Mehlis gland cell progenitors, was the only marker that was substantially up in *foxA* knockdown males and was significantly down in *foxA* knockdown females. Since *foxA* is not enriched in the Mehlis gland, we again speculate that this might be due to the variations in the reproductive status of males and females. Levels of *cbp1* were also similar between control and *foxA* knockdown worms. Together, these data suggest that FoxA is primarily restricted to the EG cell lineage, and that EG loss does not significantly perturb the balance or heterogeneity of stem cells during homeostasis, at least during the first two weeks of *in vitro* culture.

### Targeted RNAi screening using *in vitro* leukocyte feeding identifies MEG-8.2 as a key EG factor necessary for lysing ingested leukocytes

Previously, we reported that without the EG, ingested leukocytes fail to be degraded in the esophageal lumen, resulting in their accumulation in the gut (*13*). This result indicates that one or more EG factors induce rapid lysis of ingested leukocytes within the lumen. Such a function appears intimately linked to parasite survival *in vivo* since nearly all *foxA* knockdown parasites are cleared from the host vasculature. Therefore, we sought to determine if one or more EG factors would play a dominant role in lysing leukocytes. We systematically knocked down 32 EG genes individually, fed peripheral leukocytes derived from *UBC-GFP* mice (*28*) to adult worms between 2-4 hours, and quantified the number of males with more than one intact cell in the anterior gut lumen (Methods) (**Figure 3A**). We could not reliably quantify females since their range of motion is significantly greater than the males. Similar to our previous findings, while a basal level of males (∼20%) contained GFP+ cells in the gut, *foxA* knockdown males showed a significantly higher fraction of worms (∼60%) bearing GFP+ cells in the gut (**Figures 3B and 3E**). We recorded at least 20 males from two or more independent experiments. To compare the differences across each gene knockdown, we normalized each data by subtracting the percentage value of the control from each data point, bringing the control value to zero (see Methods) (**Figure 3C**). Intriguingly, we found two candidate genes with a significant increase in the fraction of males containing GFP+ cells in the gut lumen – *meg-8.2* (Smp_172180) and *ppt-1* (Smp_142980). Of the two, *ppt-1* showed greater variability, while *meg-8.2* consistently showed an increase across multiple independent experiments (**Figures 3D and 3E**). To determine if *meg-8.2* knockdown perturbs EG integrity, we first analyzed the expression changes of EG genes using qPCR (**Figure 3F**). All tested EG genes were expressed at a similar level to the control worms, suggesting that *meg-8.2* is specifically downregulated. In addition, most *meg-8.2* knockdown worms’ EG was positively labeled by PNA (**Figure 3G**). These data indicate that *meg-8.2* knockdown does not perturb EG integrity but phenocopies *foxA* knockdown, in which the parasites fail to capture and degrade ingested leukocytes in the esophagus.

**Figure 3.**
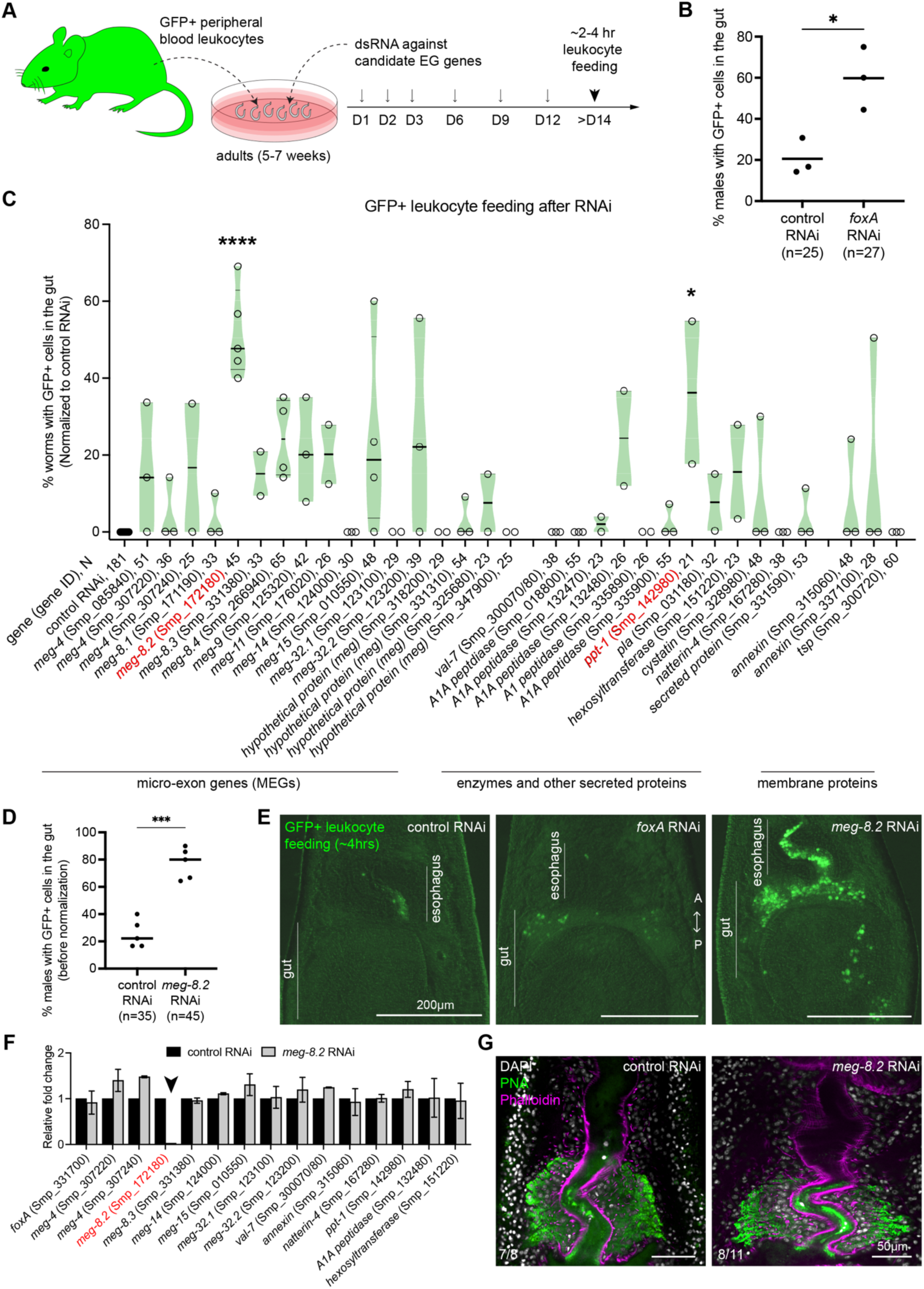
*meg-8.2* knockdown does not affect EG integrity and phenocopies *foxA* knockdown in its failure to degrade ingested leukocytes. (A) Experimental scheme to knock down individual EG genes and feed the parasites with leukocytes derived from peripheral blood of GFP-expressing mice. (B) Three independent experiments show an increased percentage of worms (males) with one or more detectable GFP+ leukocytes in the anterior gut lumen. n: total worms from three independent experiments. RNAi screen of 33 EG genes. Each dot represents the percentage of worms (males) with one or more detectable GFP+ leukocytes in the anterior gut lumen in each independent experiment. The percentage value was normalized by subtracting the percentage value of control worms in that experiment. Between 2 to 5 independent experiments were performed for each gene. N: total worms analyzed in all experiments. Ordinary one-way ANOVA was performed to analyze the statistical significance. (D) Unnormalized data for *meg-8.2* knockdown shown in (C). Paired t-test was used to determine the significance. (E) Images of the head region show an accumulation of GFP+ leukocytes in the gut lumen of *foxA* and *meg-8.2* knockdown males. (F) qPCR of EG genes in *meg-8.2* knockdown showing a specific downregulation of *meg-8.2* and relatively similar levels of other genes. cDNA samples were derived from control and *meg-8.2* knockdown adult males. (G) PNA and Phalloidin labeling shows that EG appears largely unperturbed. The numbers indicate the male worms analyzed.

### Recombinant MEG-8.2 directly lyses host leukocytes and erythrocytes

Our RNAi screen suggests that MEG-8.2 could play a dominant role in degrading ingested leukocytes, meriting a deeper investigation into its potential mechanism of action. MEG-8.2 is one of the four members of the MEG-8 family (*29*). Previous studies identified over 50 transcripts that belong to the MEG family (*15, 16, 29, 30*). To better understand the relationship between MEG-8 family genes and other MEGs, we retrieved amino acid sequences of all identifiable MEGs from the *S. mansoni* genome (SM_V10) (*15*) and analyzed their phylogeny. MEG-8.2 closely aligned with MEG-8.3 (Smp_331380) and MEG-8.4 (Smp_266940), while MEG-8.1 (Smp_171190) was more distant (**Figure S4**). As reported, when BLASTed against other members of the *Schistosomatidae* family, it was evident that closely related species such as *S. rodhaini*, *S. haematobium*, *S. japonicum*, and *S. bovis*, as well as more distant avian/animal schistosomes (e.g., *Trichobilharzia*, *Heterobilharzia*) carry several MEG-8-family orthologs (**Figure S5A**) (*29*). MEG-8 family proteins are relatively small, and all have a ∼20 amino acid N-terminal signal peptide, suggesting that this region is likely cleaved upon secretion. AlphaFold predicts that towards the C-terminus, MEG-8.2 has three helices (α1: 50 – 62; α2: 91 – 103; and α3: 115 – 138 amino acid positions) (**Figure 4A**). MEG-8.2 amino acid positions E107, P110, W121, L123, F124, F128, and L129 are conserved among the MEG-8 family proteins (**Figure S5B**). These residues span the disordered linker (L3) between α2 and α3 (104–114) and α3 (115–138). Previous studies hypothesized that MEGs are alternatively spliced to produce distinct protein variants (*15, 16*). However, the functional relevance of the potential protein variants remains unclear. *meg-8.2* consists of 17 exons, 15 of which are micro-exons ranging between 12 to 33 base pairs. To determine the predominant form of splice variants, we generated and sequenced ten independent clones from the total cDNA of adult males, adult females, and day one and day seven schistosomula. Only one out of 10 clones carried a cDNA with a skipped exon 11 (**Figure S5C**), which results in a non-frameshift mutation that lacks ten amino acids (79 – 88aa) located in the disordered linker (L2) between α1 and α2. However, the rest of the clones were at the full length.

**Figure 4.**
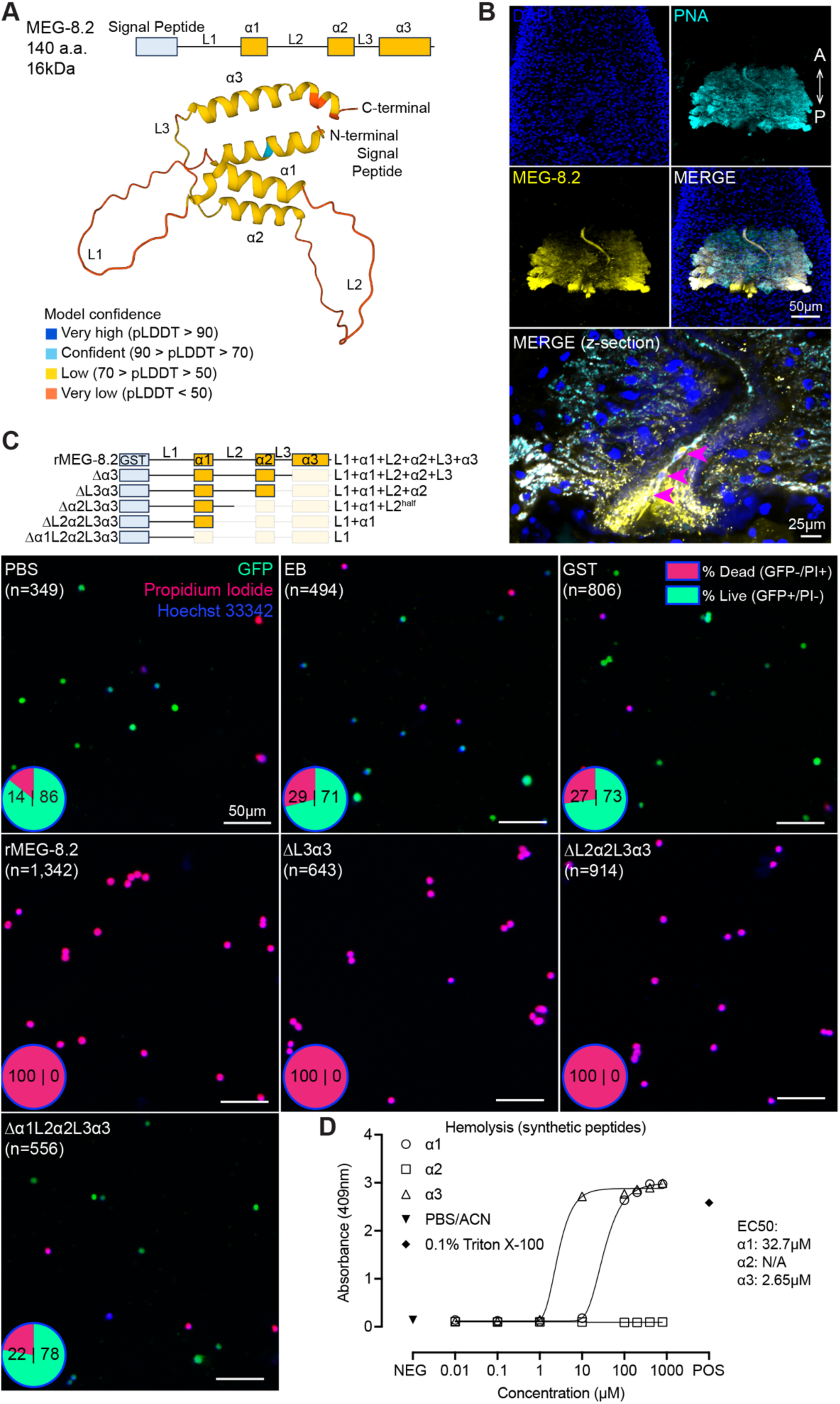
MEG-8.2 helices directly lyse host leukocytes and erythrocytes. (A) AlphaFold structure prediction shows an N-terminal signal peptide and three helices spanned by three disordered linkers. (B) Confocal image of an adult male head labeled with PNA lectin (cyan) and α-MEG-8.2 (yellow). Maximum Intensity Projection is shown for low-magnification images, and the merged high-magnification image of a different male (below) is shown as a single confocal z-section. Arrowheads show host leukocytes in the esophageal lumen. (C) GFP-expressing leukocytes were treated with truncated MEG-8.2 mutants for 10 minutes at 37°C. Each panel shows an overlaid representative image of GFP, PI (magenta), and Hoechst33342 (blue) for controls (PBS, elution buffer, GST only) and mutants. % viability is shown as a pie chart in the lower left corner of each image. The two numbers inside each pie chart are the percentages of dead (PI+/GFP-) and live (PI-/GFP+) cells. n: total number of cells counted. (D) Hemolytic activity of synthesized α1, α2, and α3 peptides. α1 and α3 peptides were dissolved in PBS/ACN and α2 in PBS.

To understand the function of MEG-8.2, we first generated custom α-MEG-8.2 antibodies against a C-terminal epitope (see Methods), which labeled most of the EG cells (**Figure 4B**). MEG-8.2 was also detected inside the esophageal lumen and surrounding the leukocytes caught in the lumen. Together with RNAi screen results (**Figure 3**), these data suggest that EG-secreted MEG-8.2 directly or indirectly degrades the ingested host leukocytes. To test this hypothesis, we replaced the N-terminal signal peptide with a GST affinity tag and expressed the full-length recombinant MEG-8.2 in *E. coli* (GST-rMEG-8.2, ∼40kDa) (**Figure S6A**) (see Methods). Upon induction of *E. coli* harboring an empty plasmid with isopropyl β-D-thiogalactopyranoside (IPTG), GST-rMEG-8.2 (referred to as rMEG-8.2 hereafter) (∼40kDa) or GST-only (26kDa) was induced at the correct size band, which was purified using immobilized glutathione beads (**Figure S6B and S6C**). For GST-rMEG-8.2, we noted several smaller size proteins that appear to be degradation products. We also noted that a portion of the induced protein was insoluble. Despite minor technical challenges, the final eluant after bead purification included full-length rMEG-8.2, confirmed by western blotting using α-MEG-8.2 antibodies.

To determine if MEG-8.2 plays a direct or an indirect role in degrading host leukocytes, we isolated GFP-expressing leukocytes from the peripheral blood of *UBC-GFP* mice, treated the cells with GST-only or rMEG-8.2 for 10 minutes at 37°C, and stained the cells with propidium iodide (PI) and Hoechst (**Figure 4C**). Cells treated with PBS, elution buffer (EB), or GST retained viability (GFP+/PI-) above 70%. Intriguingly, we observed that virtually all cells treated with rMEG-8.2 had lost cytoplasmic GFP signal and stained positively with PI, indicating that MEG-8.2 can directly lyse the leukocytes. In addition, rMEG-8.2, but not GST-only or EB control, also displayed activity against the whole peripheral blood, indicating that MEG-8.2 is also capable of lysing erythrocytes (**Figure S6F**). To determine if other MEG-8 proteins display similar activity against host cells, we recombinantly purified MEG-8.1, MEG-8.3, and MEG-8.4 using the same approach (**Figure S6D**) and tested them on leukocytes and erythrocytes (**Figures S6F and S6G**). Surprisingly, while rMEG-8.1 and rMEG-8.4 did not lyse the cells, rMEG-8.3 also induced lysis of leukocytes and erythrocytes. Sequence alignment indicates that MEG-8.2 has more shared residues with MEG-8.3 than with MEG-8.1 or MEG-8.4 (**Figure S5B**). Together, these results support a ‘direct’ role of MEG-8.2 in lysing ingested leukocytes within the esophageal lumen.

How does MEG-8.2 lyse host cells? As reported, similar to helices of other MEGs (*14, 29, 31*), the three predicted MEG-8.2 helices have a hydrophobicity index between 0.4 and 0.8, with several non-polar residues biased towards one-half of the helical surface, indicating their likely amphipathic nature (**Figure S6H**). To determine if any of the three helices contribute to its activity against host blood cells, we generated a series of truncation mutations from the C-terminus and purified the proteins (**Figure 4C, S6E, and S6G**). Surprisingly, rMEG-8.2 mutants lacking α3, L3, α2, and L2 all retained the lytic activity against the isolated leukocytes, suggesting that these domains are dispensable for this activity. The lytic activity was lost only when the deletion included the α1 domain, suggesting that α1 is sufficient and is the minimum domain required to induce cell lysis. Similar results were observed in the hemolysis assay (**Figure S6F**), indicating that MEG-8.2 α1 might also be directly lysing erythrocytes.

To determine if the lytic activity of MEG-8.2 helices is concentration-dependent, we synthesized the three peptides (α1: FWRRMWNSFTSMF; α2: LKERIMNKFNSIF; α3: FTERLWMLFKHCFLNFKNLA-KIF) and tested a range of concentrations using a hemolysis assay (**Figure 4D and S6I**). While α2 did not lyse the cells, α1 and α3 displayed hemolytic activity at micromolar concentrations (EC_50_: 32.7 – 60.8µM and 2.7 – 7.8µM, respectively). Surprisingly, although α3 displayed a higher activity than α1, it was dispensable for lysis from our assays using recombinant proteins. Together, these results suggest that while α1 is sufficient for cell lysis, α3 might enhance the lytic activity.

### MEG-8.2 interacts with host cytoplasmic and membrane proteins

Having identified a key EG factor that blocks and degrades ingested host cells, we sought to tease apart whether such a function is linked to the development and survival of schistosomes *in vivo*. Previously, we showed that when transplanted into a naïve host, adult schistosomes lacking the EG (via *foxA* knockdown) can survive for a week after transplantation (*13*). However, their survival drops significantly between 2-4 weeks post-transplantation, and a few remaining survivors are severely stunted. To determine if EG-lacking schistosomes have defects during the development of the parasite and whether the MEG-8.2-driven blood cell lysis impacts the development and survival of juveniles, we mechanically transformed cercariae to schistosomula, divided them evenly into three groups, treated them with control, *foxA* or *meg-8.2* dsRNA for a week, intravenously injected an approximately equal number of schistosomula, and collected the parasites after two weeks via hepatic perfusion (see Methods) (**Figure 5**). *foxA* knockdown juveniles were significantly shorter either due to the larger worms (that consume more blood) being cleared by the host and/or due to an unknown EG/FoxA-dependent developmental mechanism. We note that while this method is limited in quantitatively assessing the survival rate due to the inherent technical variability (e.g., loading an equal number of parasites, injecting the parasites into the vein), *foxA* knockdown parasites were generally recovered at a lower number compared to controls. In contrast, *meg-8.2* knockdown juveniles showed a similar size distribution to control parasites, suggesting that other EG factors might play additional roles and that the EG-mediated immune evasion mechanism likely encompasses a broader scope beyond the initial steps of lysing the ingested leukocytes.

**Figure 5.**
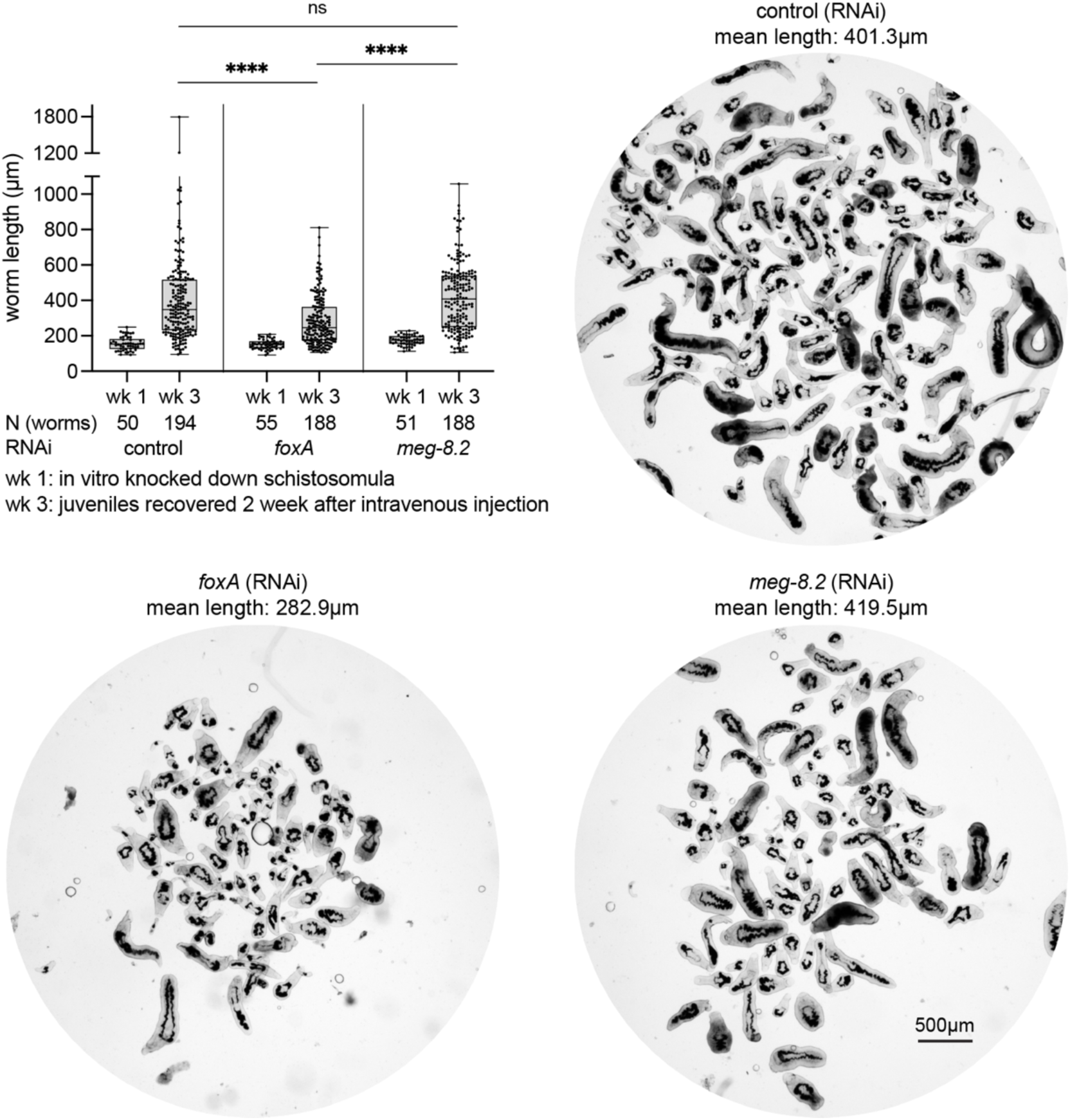
*meg-8.2* knockdown does not perturb *in vivo* parasite development and survival. Juvenile parasites were recovered two weeks after intravenous injection of schistosomula treated *in vitro* with dsRNA for a week. While *foxA* RNAi juveniles are significantly shorter, *meg-8.2* RNAi juveniles have a similar size distribution as control parasites. One-way ANOVA (Tukey’s multiple comparison test) was used for statistical analysis. N: total number of worms analyzed for length measures. The mean length value for each juvenile group is indicated above each image.

Why do schistosomes secrete large amounts of MEG-8.2 into the lumen to degrade incoming host cells if the ingested cells are likely eventually degraded in the gut lumen, and their degradation by MEG-8.2 appears not essential to the survival of the parasites? We speculated that MEG-8.2 might have additional roles beyond the local lysis of ingested blood cells for two reasons. First, although the individual helices have high hydrophobicity and hydrophobic moment, they are relatively a small part (less than 50%) of the entire sequence – the residues between the helices appear disordered. These regions might assist the helices in interacting with membrane lipids or have a separate role in protein-protein interactions. Such a possibility is substantiated by a previous report of another EG factor (e.g., MEG-14) that was shown to interact with an immunomodulatory protein S100A9 from a yeast two-hybrid screen (*32*). Second, known amphipathic helices in other animals play diverse roles, from antimicrobials (*33*) to receptor binding and signal transduction (*34*). Thus, we hypothesized that MEG-8.2 may have a role in parasite-host protein-protein interactions. To identify potential interacting proteins, we used rMEG-8.2 to pull down whole-blood lysate and performed LC-MS/MS (see Methods) (**Figure 6A and Table S2**). Excluding false positives by crosschecking any shared hits with the negative controls (blank, prey only, and GST pull-down), we identified 203 hits (UniProt accession IDs), of which 30 had an overall score above zero with two or more unique peptides (**Figures 6A and 6B**). The top four candidates among them were Cct2 [P80314], Memo1 [Q91VH6], Cd98hc [P10852], and Ltf [P08071]. To confirm the specificity of these interactions, we performed Western blots of the top three (i.e., Cct2, Memo1, and Cd98hc) proteins in GST-only and rMEG-8.2 pull-down samples (**Figure 6E**). We observed bands corresponding to the expected size of the host proteins in the rMEG-8.2 pull-down but not in the GST-only pull-down, suggesting that these are true positive interactions.

**Figure 6.**
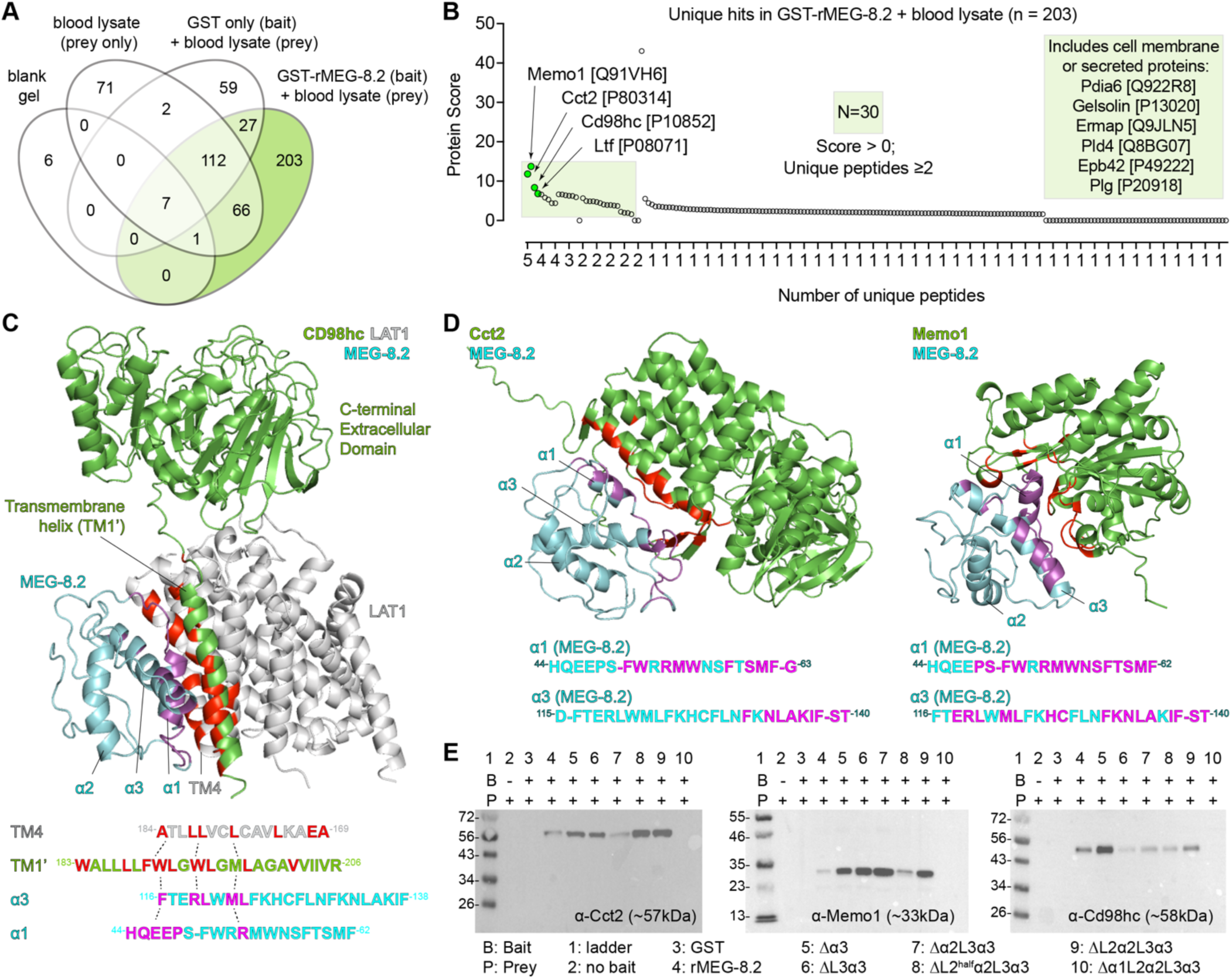
MEG-8.2 interacts with host blood cell proteins. (A) Venn diagram summary shows the number of hits uniquely identified in GST-rMGE-8.2 pull-down (**Table S2**). (B) Dot plot showing the overall score and the number of unique peptides of the 203 hits. The top four candidates are highlighted with green dots. The light green square includes 30 hits with an overall score above 0 and two or more unique peptides. (C-D) Interaction modeling between MEG-8.2 and CD98hc-LAT1 complex (*35*) is shown in (C), while MEG-8.2-Cct2 and MEG-8.2-Memo1 models are shown in (D). MEG-8.2 residues with an interaction distance <4Å are colored in magenta. Host proteins are labeled in green except for LAT1 (gray), which is not part of the mass spectrometry hits. The counterpart residues of the host protein that interact with MEG-8.2 residues are highlighted in red. (E) Western blots of Cct-2, Memo1, and CD98hc in prey (blood lysate) only (lane 1), pull-down samples using GST only (lane 2), or pull-down samples using rMEG-8.2 full lengths and deletion mutants (lanes 3 to 10).

To determine if any of the helices or the disordered regions are required for the interactions, we first took an *in silico* approach by modeling the MEG-8.2 interaction with the candidate proteins (see Methods) (**Figures 6C, 6D, and S7**). For example, using published structural data (*35*), we modeled MEG-8.2 with CD98hc (also known as Slc3a2), a transmembrane protein that is reported to form a heterodimer with Slc7 family transporters and functions as an amino acid transporter in integrin signaling and adaptive immunity (*36–39*). Interestingly, several hydrophobic residues of α1 and α3 showed close interaction with the hydrophobic residues of the CD98hc transmembrane helix (TM1’) (**Figure 6C**). Hydrophobic residues of TM1’ (F189, W193, M196, L197, A200, I203) that normally interact with hydrophobic residues of LAT1 (also known as SLC7A5) TM4 (L173, L177, L181) are closely positioned with the hydrophobic residues of MEG-8.2 (e.g., α3 F116, L120, M122, L123), suggesting that MEG-8.2α1/α3 -TM1’ interaction might interrupt the CD98hc-LAT1 interaction. Similarly, hydrophobic residues of α1 and α3 showed close interactions with Cct2 and Memo1 (**Figure 6D**) as well as Ltf (Lactotransferrin) (**Figure S7A**), a known component of neutrophil secretory granule, with antimicrobial properties (*40–42*). Interestingly, the western blot of host proteins after a pull-down with the truncation mutants revealed that α3 was dispensable for these interactions, while α1 was necessary (**Figure 6E**). Furthermore, protein lysate from the plasma of the infected host contained MEG-8.2 but not of the uninfected host (**Figure S7B**) suggesting that MEG-8.2 is likely released into the blood. These results suggest that MEG-8.2 α1 plays a dual role: concentration-dependent lysis of the host blood cell membrane and interaction with host proteins related to the proper function of immune defense. This proposes an interesting possibility that at a high (i.e., cell lysing) concentration inside the EG lumen, MEG-8.2 contributes to the effective degradation of incoming blood cells. In contrast, MEG-8.2 that is released into the host bloodstream (presumably via worms’ regurgitant) is at a low concentration (that cannot lyse cells) but interacts with host cell membrane or secreted/excretory proteins, thereby inhibiting the signaling and activation of the immune system (**Figure 7**).

**Figure 7.**
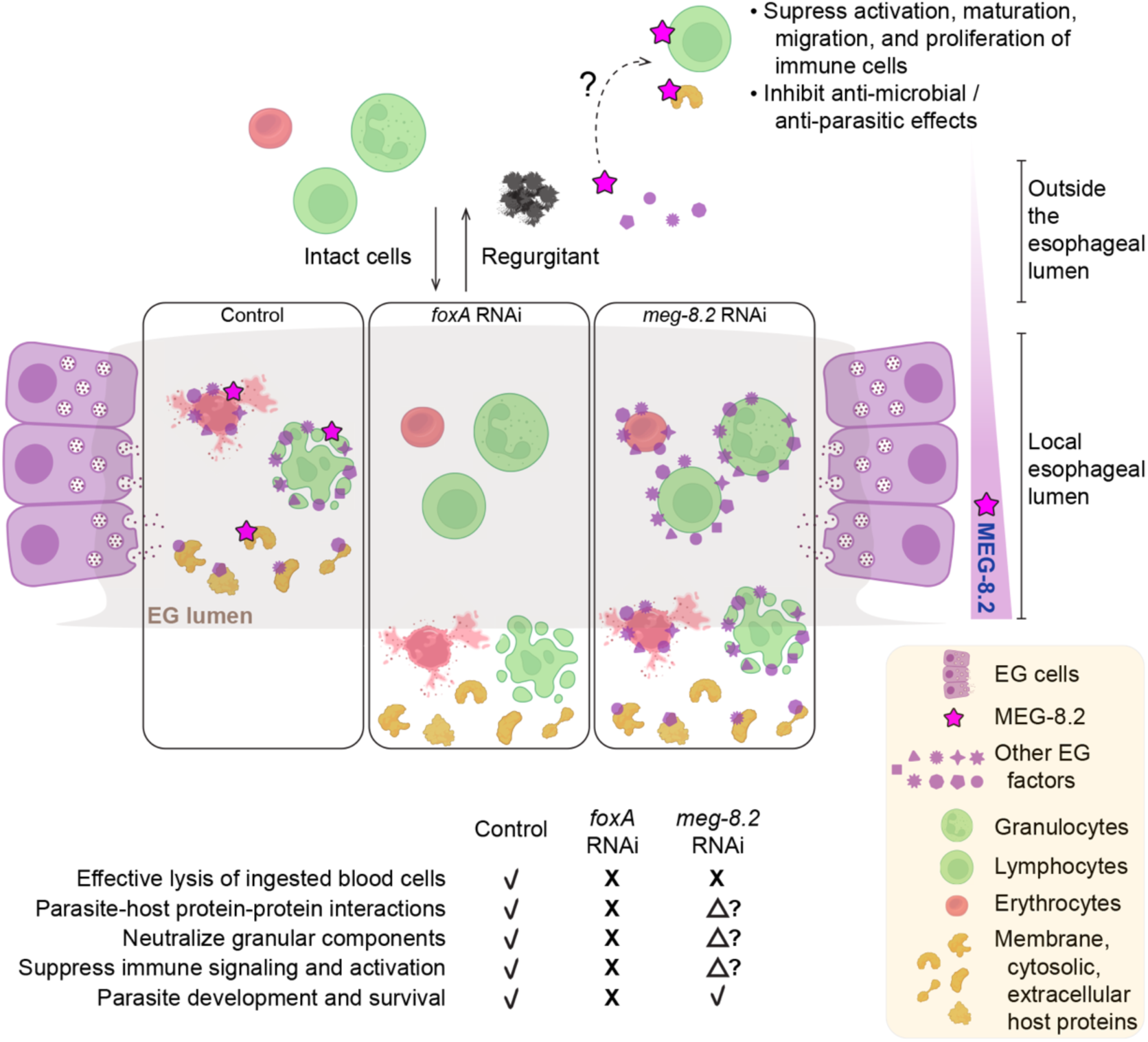
Working model for MEG-8.2-mediated host immune evasion. Model for MEG-8.2-mediated host immune evasion. In control parasites, released EG factors effectively lyse the ingested host cells in the lumen and interact with host proteins that promote parasite survival. MEG-8.2 may also be released into the bloodstream via worms’ regurgitation to interact with host extracellular and membrane proteins to suppress the activation of immune cells. In the absence of EG, no EG factors exist to lyse the cells or interact with host proteins, leading to perturbed parasite development, homeostasis, and survival. In MEG-8.2 absence, host cells are inefficiently degraded, and interactions between other EG factors and host proteins are largely unaffected. The model was generated using BioRender and Illustrator (Adobe).

## Discussion

As schistosomes ingest large amounts of blood during growth and homeostasis, diverse host cells and proteins enter the digestive tract. These host components can presumably be digested when neutralized and used as a potential nutritional source for the parasites. If not, many of the components could potentially damage the parasite tissue, either directly by releasing proteases and peroxidases (e.g., granulocyte secretion) that could be damaging, or indirectly by relaying the signals to activate, mobilize, and recruit more host immune factors and/or cells. Thus, as the frontline defense, the EG must not only degrade incoming host cells but also neutralize any proteins that could harm the parasites. EG factors have been gaining attention over the past two decades; however, the complete picture of the EG-expressed genes and their functions has remained elusive. Having discovered the essentiality of FoxA in EG cell production (*13*), using a comparative transcriptomics approach, we were able to identify known and new EG factors. The identified factors are expressed in most, if not all, EG cells without substantial heterogeneity, and most of them are shared between male and female schistosomes.

From a targeted RNAi screen, we discovered that depletion of MEG-8.2 leads to an accumulation of leukocytes in the gut. While previous studies have hypothesized that MEGs could confuse the host immune system by producing protein variants through exon skipping (*12, 16*), we find that most MEG-8.2 might exist in full length. We note that since our functional screen analyzes a short window of time after initiating feeding, it may not identify all the candidates that have a role in lysing the ingested leukocytes. Indeed, while MEG-8.2 displayed the most severe phenotype in our assay, other EG factors showed moderate phenotypes, including several MEGs (e.g., MEG-4, MEG-8.3, MEG-8.4, MEG-9, MEG-11, MEG-15, and MEG-32.2) and enzymes (e.g., A1A Peptidase, Ppt-1, and Hexosyltransferase). We also note that the screening pipeline does not pick up parasites with altered physiological or cellular changes associated with each gene knockdown. Recent work suggests that *meg-8.3* knockdown shows EG degeneration over time, rendering parasites sick (*43*). Although in our hands most of the *meg-8.3* knockdown worms remained attached to the culture dish at the time of feeding, it is possible that the worms did not ingest leukocytes as much as *meg-8.2* knockdowns. However, a single gene knockdown of *meg-8.2* that results in leukocyte gut accumulation in a significant proportion of worms suggests that MEG-8.2 might be the dominant driver of this process.

Our biochemical investigations using recombinant MEG-8.2, as well as synthetic peptides, reveal the crucial role of individual helices in lysing host leukocytes and erythrocytes. While α1 is the minimal region needed for this activity, α3 is ∼10 times more potent, and thus, the two domains likely work together to efficiently lyse host cells. Both helices have amphipathic properties (**Figure S6H**). Amphipathic helices are found in diverse aspects of biological processes that play significant roles in membrane curvature sensing, remodeling, and permeabilization (*44–46*). Thus, the hydrophobic side of MEG-8.2 α-helices likely inserts between the acyl side chains lipid bilayer of the host cell membrane, altering the membrane curvature, resulting in permeabilization. In addition, as reported for other MEGs (e.g., MEG-14) (*47*), MEG-8.2 linkers that connect the helices appear disordered (**Figure 4A**). These internally disordered regions fold differently in varying environmental conditions such as increased temperature, presence of negatively charged lipids/detergents, and dehydration, suggesting that they may contribute to the context-dependent roles of MEG-8.2 in targeting different cells and molecules. Meanwhile, previous studies report that secondary structure prediction of several MEGs (e.g., MEG-9, −12, −19, −24, −26, −27, and −28) reveal that they also have amphipathic helices (*14, 29, 31*). This is consistent with the fact that MEG-8.3 has a C-terminus with a high sequence similarity to MEG-8.2 α3 (**Figure S5B**), which is likely responsible for its lytic activity (**Figures S6F and S6G**). Given that several MEGs show a moderate increase in leukocyte gut accumulation from our functional screen (**Figure 3C**), it will be important to tease apart the relative contributions of amphipathic helices of other MEGs in degrading ingested blood cells.

The fact that depletion of MEG-8.2 alone does not appear to have a major impact on parasite development or survival but that it interacts with multiple host proteins has important implications. Cell lysis by the EG factors is likely one of the first steps in neutralizing host components. For instance, permeabilized granulocytes may release several factors, such as proteases and peroxidases packed in cytotoxic granules. Neutralizing or sequestering these components in a timely fashion would minimize damage to the parasites. We speculate that although MEG-8.2 is the dominant driver of host cell lysis in the esophageal lumen, different EG factors interact with distinct host proteins to inhibit their activity before sending them into the gut. This model is supported by the fact that MEG-14, another prominent EG factor, is shown to interact with S100A9 (*32*), a calgranulin protein expressed in granulocytes (e.g., neutrophils) that promotes and regulates inflammatory processes (*48, 49*). According to this model (**Figure 7**), in *foxA* knockdown, all EG factors are absent, and therefore, the parasites likely accumulate damage over time and can no longer survive. However, other EG factors are still present in *meg-8.2* knockdown (such as MEG-8.3, given its recombinant form can also lyse host cells), which is why parasites are not affected. This model also explains why other attempts to inhibit a single EG factor (e.g., MEG-4) have not resulted in a dramatic increase in parasite clearance (*50*).

Our comparative mass spectrometry of rMEG-8.2 pull-down reveals that it interacts with various host membrane, cytosolic, and extracellular proteins. The top two hits, Cct2 and Memo1, are cytoplasmic proteins. Cct2 (T-complex protein 1β), a component of the chaperonin-containing T-complex (TRiC), is involved in protein folding and proteostasis (*51, 52*) by acting as an aggrephagy receptor, binding to misfolded aggregation-prone proteins for clearance (*53*). Memo1 (mediator of ErbB2-driven cell motility 1) promotes cell proliferation and migration (*54*), and recent evidence suggests that it binds to Copper (Cu) and serves as a redox enzyme to sustain O_2_^−^ production by NADPH (*55, 56*). Previous studies report their presence in extracellular vesicles (*57–61*), suggesting that their interaction with MEG-8.2 may occur outside of the host cell. Whether these interactions occur inside (e.g., MEG-8.2 being phagocytosed or entering the permeabilized cell’s cytoplasm) or outside (e.g., permeabilized cells releasing cytosolic contents into the lumen or released via extracellular vesicles) the host cell remains to be determined. In either case, MEG-8.2’s interaction with these proteins may have functional relevance that is advantageous to the parasites (e.g., preventing protein aggregation or lowering oxidative stress). CD98hc (amino acid transporter heavy chain SLC3A2) is a membrane protein with a single transmembrane (TM1’) helix and a C-terminal extracellular domain. As modeled (**Figure 6C**), MEG-8.2 α1 and α3 preferentially interact with TM1’ via hydrophobic interactions with residues that are important in forming a heterodimer with SLC7 protein (*35*). CD98hc facilitates the integrin-dependent proliferation of T and B lymphocytes that shape adaptive immunity (*39, 62*). MEG-8.2 presence in blood plasma suggests that it is excreted from parasites. The excreted MEG-8.2 (at low concentration) may interact with live CD98hc-expressing T and B lymphocytes (in the bloodstream or reaching further to the peripheral lymphoid organs) to inhibit their activation/proliferation (**Figure 7**). Lactotransferrin (Ltf) binds to iron and is found in serum and secretory fluids (e.g., breast milk, saliva) and secretory granules of neutrophils (*40, 42, 63*). It also has antiviral, antimicrobial, antifungal, and antiparasitic activity (*64, 65*). Furthermore, Ltf binds to different receptors (e.g., lipoprotein-related receptor) and is internalized, which activates innate and adaptive immune cells (*66*). Similarly, further down the list of MEG-8.2 interacting proteins is Gelsolin (**Table S2**), which is another neutrophil granular component (*42*). These results suggest that MEG-8.2 potentially interacts with granular proteins via hydrophobic interactions of α1 (and α3) to prevent them from damaging the parasites and inhibit their signaling and activation of innate and adaptive immunity.

Further investigations are necessary to dissect the potential role and the mechanism of MEG-8.2-host protein interactions in suppressing host immunity. In addition, devising novel *in vitro* approaches to identify the role of EG factors other than lysing the ingested leukocytes will be crucial in broadening our understanding of their functional relevance to host immune evasion. A recent study using epitope mapping of proteins expressed in the digestive tract and tegument reveals potential in designing multi-epitope vaccine constructs that can serve as monoclonal antibody targets (*67, 68*). Interestingly, several EG MEGs are considered among the top candidates, suggesting their therapeutic potential. In this regard, uncovering the host interaction partners of other EG factors and identifying a combination that causes a significant *in vivo* parasite clearance will have important implications in devising new approaches to target schistosomes.

## Methods

### Animal care and handling

Vertebrate animals were handled in accordance with the Institutional Animal Care Use Committee protocols at the University of Wisconsin–Madison (M005569) and The University of Texas Health Science Center at Houston (AWC-21-0069 and AWC-24-0073). Swiss-Webster mice (Hsd:ND4, Envigo or SW-F, Taconic Biosciences) between 3 to 5 weeks of age were infected with *S. mansoni* (NMRI strain) received from the Biomedical Research Institute (Rockville, MD). Infected mice were harvested via hepatic perfusion (*69*) between 5 to 7 weeks to collect adult parasites. Peripheral blood was collected from Swiss-Webster, C57BL/6J (#000664, The Jackson Laboratory), or *UBC-GFP* (#004353, C57BL/6-Tg(UBC-GFP)30Scha/J, The Jackson Laboratory). For infection with schistosomula, female Swiss-Webster mice, age three to five weeks were used. Mice were weighed and randomly assigned to three treatment groups until infection. Prior to infection, mechanically transformed schistosomula were treated *in vitro* with 15ng/µL of control, *foxA*, and *meg-8.2* dsRNA five times for a week. RNAi schistosomula were washed three times in 1 x PBS (Calcium and Magnesium free) and ∼3,000 larvae were loaded into 27G insulin syringe (BD) in ∼200µL maximum volume. Following anesthesia using isoflurane (6679401725, Piramal Critical Care), mice were placed laterally and schistosomula were injected retro-orbitally after conforming aspiration of needle placement in the retro-orbital sinus. Post-inoculation, mice were housed in a group and were monitored daily. Mice were euthanized three weeks post-infection using Fatal-Plus (Vortech) / Heparin (McKesson) cocktail and worms were recovered via hepatic perfusion.

### RNA interference

Previously published methods were used to synthesize double-stranded RNA for all genes reported in this study (*13, 70*). **Table S3** lists all oligonucleotides used for dsRNA synthesis. Briefly, ∼15 to 20 adult parasites were cultured in ABC media (*71*) without the red blood cells for ∼2 weeks *in vitro*. Parasites were treated with dsRNA between 10 – 20µg/mL 6 times during the culture period. Knockdown adult worms were used for RNA-seq, qPCR, staining, and leukocyte feeding. For schistosomula knockdown, mechanically transformed schistosomula (*69, 72*) were split equally into three conditions (control, *foxA*, *meg-8.2*) and cultured for a week while being treated with dsRNA 4 to 5 times. Knockdown schistosomula were injected retro-orbitally and juveniles were collected 2 to 3 weeks after the injection.

### Quantitative real-time PCR

Total RNAs from knockdown parasites were extracted using TRIzol (15596026, Invitrogen)/phenol-chloroform. Extracted RNA samples were converted to cDNA using iScript cDNA synthesis kit (Bio-Rad). qPCR was performed using CFX Real-Time PCR System (Bio-Rad). Relative fold changes were analyzed using ΔΔCt method. Primers used for qPCR are listed in **Table S3**.

### RNA-seq analysis

RNA sequencing (2 x 150bp, ∼30 million reads per sample) was performed at the University of Wisconsin Biotechnology Center (Madison, WI). Sequenced reads were processed and analyzed using CLC Genomics Workbench (Qiagen) under default settings. Quality passed reads were mapped to the *S. mansoni* genome (SM_V9 assembly). Differential expression analysis was performed to determine statistical significance (false discovery rate ≤ 0.05; absolute Log_2_ fold change ≥ 1). Raw and processed reads have been deposited in NCBI (GSE278682).

### Parasite labeling and imaging

Colorimetric and fluorescent *in situ* hybridizations were carried out using previously described methods (*13, 19, 26, 70, 73, 74*). Briefly, adult parasites were treated with 2.5% tricane (ethyl 3-aminobenzoate methanesulfonate) to separate the males and females and killed with 0.6M MgCl_2_ before fixing in 4% formaldehyde/PBSTx for ∼4 hrs. Fixed worms were dehydrated in methanol and stored at −20°C. Rehydrated worms were bleached in 0.5% formamide/1.2% H_2_O_2_/0.5x SSC for 45 – 60 minutes under bright light and treated with 10µg/mL Proteinase K (Invitrogen)/0.1% SDS for 30 minutes. Riboprobes were synthesized by *in vitro* transcription. Candidate gene fragments inserted into pJC53.2 were amplified by PCR and were used as a template for a transcription reaction containing either Digoxigenin-11-UTP (DIGUTP-RO, Roche) or Fluorescein-12-UTP (11427857910, Roche) and either T3 or SP6 polymerase, depending on the orientation of the insert, to generate antisense riboprobes. Purified riboprobes were hybridized to parasites at 52 °C overnight. Worms were incubated with anti–DIG-AP (11093274910, MilliporeSigma) for colorimetric, and anti-DIG-POD (11207733910, MilliporeSigma) or anti–FITC–POD (11426346910, MilliporeSigma) for fluorescent in situ hybridization at 1:1,000 – 1:2,000 dilution overnight. Peanut Agglutinin tagged with fluorescein (VectorLabs) was used at 1:500 to label the esophageal gland. Polyclonal anti-MEG-8.2 antibodies were custom ordered commercially (ThermoFisher Scientific) by injecting a short C-terminal peptide (EEYNPPKDSDFTER) into a rabbit host. Affinity-purified anti-MEG-8.2 antibodies were used at 1:500 for immunofluorescence staining in FISH blocking solution. Secondary goat anti-rabbit IgG (H+L) tagged with Alexa Fluor 568 (A-11036, ThermoFisher Scientific) was used at 1:500 in FISH blocking solution. Low-resolution colorimetric and fluorescent images were taken using Leica M205FCA stereoscope or AxioZoom.V16 stereomicroscope (Carl Zeiss). For high-resolution images, Andor WDb spinning disk confocal microscope (Andor Technology) or Nikon A1 confocal microscope. ImageJ or ImarisViewer (Bitplane) was used to make linear adjustments to brightness and contrast. Schistosomula and juvenile RNAi worms were imaged using ECHO Revolve (BICO).

### In vitro leukocyte feeding

∼15 – 20 adult parasites were knocked down for individual EG genes for two weeks *in vitro*. Isolated total leukocytes from one to two *UBC-GFP* mice were resuspended in 0.5 – 1 mL of cold 1x PBS. Between 50 – 150µL of leukocyte suspension was added to each knockdown. The volume of cells added to each well differed slightly across each experiment (due to the differences in the total number of gene knockdowns in a particular experiment). However, within each experiment, an equal volume of cells was added to each well. After two to four hours of incubation at 37°C/5%CO_2_, parasites were treated with 2.5 % Tricane (ethyl 3-aminobenzoate methanesulfonate, E10521, MilliporeSigma) to separate the males from females and to paralyze them for imaging. Male parasites were placed on a glass slide, and a cover glass was placed to flatten the worms and to prevent drying. The anterior portion of the male worms was imaged using a stereoscope (Leica Microsystems). Parasites with at least one detectable GFP+ cell in the gut lumen were counted towards a positive phenotype (i.e., defective lysis in the esophagus).

### Motif analysis

To investigate if FoxA binds to upstream regions of the identified EG genes, we used the CentriMo (v5.5.5) tool of the MEME Suite (https://meme-suite.org/meme/index.html) (*21*). We extracted and compiled the sequences of 5,000bp upstream of the transcription start site of all 36 EG genes and used them as the input for the CentriMo analysis to identify motifs enriched in these sequences. Command lines: centrimo --oc . --verbosity 1 --local --score 5.0 --ethresh 10.0 --bfile sequences.fa.bg sequences.fa motif_db/JASPAR/JASPAR2022_CORE_redundant_v2.meme.

### Cloning, expression, and purification of recombinant proteins

Full-length rMEG-8.2 and its deletion mutants, as well as other MEG-8 proteins (i.e., MEG-8.1, MEG-8.3, MEG-8.4), were amplified by PCR from *S. mansoni* cDNA. Primers included *BamHI* and *XhoI* restriction sites at 5’ and 3’ ends, respectively. The PCR products and pGEX4T-3 expression vector were digested with *BamHI* and *XhoI*, and the PCR products were purified using a gel DNA recovery kit (D4007, Zymo Research). The digested vector and PCR products were ligated using T4 DNA ligase (M0202L, NEB). The ligation mixtures were transformed into *E. coli* DH5α competent cells. Positive clones were selected on LB agar plates containing ampicillin (100 µg/mL). Plasmid DNA was isolated using a plasmid mini-prep kit (D4020, Zymo Research) and confirmed by restriction digestion and plasmid sequencing (Plasmidsaurus). Verified constructs were transformed into *E. coli* BL21(DE3) cells for protein expression. The plasmid sequences for all constructs used for recombinant protein expression are listed in **Table S4**. A single colony from the transformation was used to inoculate 10 mL of LB broth containing ampicillin (100 µg/mL), which was grown overnight at 37°C with shaking. This culture was used to inoculate 500 mL of LB broth with ampicillin (100 µg/mL), and the culture was grown at 37°C until the OD600 reached 0.6-0.8. Protein expression was induced with 0.1 mM IPTG, and the culture was incubated at room temperature for 5 hours. Cells were harvested by centrifugation at 5000 x g for 10 minutes at 4°C. The cell pellet was resuspended in GST-fusion protein lysis buffer (0.1% Triton X-100 in PBS with 1x protease inhibitor). The suspension was sonicated on ice using a large-tip probe (1 min 40 sec total sonication time; 5 sec ON, 10 sec OFF; amplitude at 30%). The lysate was cleared by centrifugation at 20,000 x g for 10 minutes at 4°C, and the supernatant containing the soluble GST-tagged protein was collected. The supernatant was incubated with Glutathione Sepharose 4B beads (17075601, Cytiva) at 4°C for 30 minutes with gentle rotation. The beads were washed three times with GST high wash buffer (0.5 M NaCl, 0.1% Triton X-100 in PBS) and two times with GST lysis buffer. The GST-tagged protein was eluted using elution buffer (50 mM Tris-HCl, pH 8.0, 10 mM reduced glutathione). Protein concentration was determined using the BCA assay kit (23227, Thermo Scientific). Fractions containing the target protein were concentrated and buffer-exchanged into PBS to remove residual glutathione using Amicon columns (UFC501096, Millipore Sigma)

### SDS Gel electrophoresis and Western blotting

Cleaned and concentrated protein fractions were boiled for 10 minutes in 5x SDS-PAGE sampling buffer. The eluted proteins were separated by SDS-PAGE and stained with Coomassie dye for visualization. For western blot analysis, 20 µg of protein samples were loaded onto a gel and transferred onto a PVDF membrane after electrophoresis. The membranes were blocked for 2 hours at room temperature in 1% blocking solution (1% Western Blocking Reagent (WESTBL-RO, Roche) in TBS) and followed by an overnight incubation at 4°C with anti-MEG 8.2 polyclonal antibody (1:2000 dilution). The next day, the membranes were washed three times for 10 minutes each with TBS-Tween wash buffer, then incubated with HRP-conjugated anti-rabbit IgG (32460, Invitrogen) for 1 hour at room temperature. After three additional 15-minute washes, the membranes were developed using the ECL-2 Western Blotting Substrate (80196, Thermo Scientific) and visualized by ChemiDoc MP Imaging System (Bio-Rad).

### Leukocyte lysis

Peripheral blood collected from *UBC-GFP* mice was centrifuged at 500 x g for 5 minutes at 4°C. For each independent experiment, one to two *UBC-GFP* mice were used. The supernatant was removed, and 3 mL of ACK lysis buffer (A1049L-01, Gibco) was added for 3 minutes (room temperature) to lyse the red blood cells. After lysis, the cells were pelleted by centrifugation at 500 x g for 5 minutes at 4°C. The pellet was washed with cold 1X PBS, and the lysis and wash steps were repeated. After the final wash, the cells were resuspended in 500 µL of 1X PBS. Recombinant MEG-8 proteins (full-lengths and mutants) were incubated with leukocytes at a concentration of 0.5 µg/µL in a total reaction volume of 20 µL in a 96-well microplate. Negative controls, including pGEX4T-3 vector-only protein (GST only) and elution buffer only, were used. The cells were incubated for 10 minutes at 37°C. Following incubation, the cells were stained with Hoechst (50 µg/mL) and propidium iodide (1 mg/mL) and imaged using M205FCA stereoscope (Leica microsystems).

### Hemolysis

Blood collected from C57BL/6 mice was centrifuged for 5 minutes (500g, 4°C). After the centrifugation, the supernatant was removed, and packed blood cells were washed twice with 10 mL of cold 1x PBS. The cells were then diluted to 1% in 1x PBS, and 75 µL was aliquoted into a 96-well microplate. Controls included pGEX4T-3 vector-only protein (GST, negative control), PBS (negative control), and 0.1% Triton X-100 solution (positive control). MEG-8 full-length and mutant proteins were diluted to 0.5µg/µL in PBS, bringing the total reaction volume to 150 µL. The plate was incubated at 37°C for 10 minutes, then centrifuged at 500 x g for 5 minutes at 4°C. The supernatant was separated and transferred to a new well. Hemoglobin was detected in the supernatant by measuring absorbance at 409 nm using a plate reader (BioTek). Synthetic MEG-8.2 peptides were custom-ordered (Biomatik) at 95% purity. Lyophilized powders were dissolved in DMSO, 30% acetonitrile (ACN), or PBS, depending on their solubility. Peptides were added at concentrations ranging from 0.01 µM to 800 µM. 30% ACN/1X PBS or 1X PBS were used as negative controls, and 0.1% Triton X-100 as a positive control.

### Pull-down

A Pierce GST Protein Interaction Pull-Down Kit (21516, ThermoScientific) was used for the pull-down assay. 100–150µg of the full-length MEG-8.2 and the mutants were used as bait proteins. Blood lysates from the peripheral blood of infected mice were used as prey proteins. For negative controls, pGEX4T-3 vector-only protein (i.e., GST-only) was used as the bait protein, while blood lysates without a bait served as negative control. The pull-down samples were processed following the kit protocol, and the protein elutes were quantified using a BCA Assay. Subsequently, 40 µg of each sample was loaded onto SDS-PAGE gels, which were stained using silver staining. The staining procedure involved fixing the gels in a solution of 40% ethanol and 10% acetic acid for 1 hour, followed by four washes with deionized water (ddH_2_O) for 20 minutes. The gels were then sensitized with 0.02% sodium thiosulfate for 1 minute, washed three times with ddH_2_O for 20 seconds each, incubated in cold 0.1% silver nitrate solution (0.1% AgNO_3_, 0.02% formaldehyde) for 20 minutes at 4°C, and washed again with ddH_2_O for 20 seconds three times and once for 1 minute. The development was carried out in a solution of 3% sodium carbonate and 0.05% formaldehyde (*75*). The stained gel was imaged by ChemiDoc MP Imaging System (Bio-Rad). The samples were resolved on SDS-PAGE and stained with Coomassie Blue for mass spectrometry analysis. After thorough destaining, bands of interest and an equivalent background region from the gel were excised. The excised gel slices were washed twice with 50% acetonitrile in water to remove excess stain. The samples were submitted to the Clinical and Translational Proteomics Service Center at the University of Texas Health Science Center for analysis.

### LC-MS/MS

The gel band samples were subjected to In-gel digestion (*76*). An aliquot of the tryptic digest (in 2% acetonitrile/0.1% formic acid in water) was analyzed by LC/MS/MS on an Orbitrap Fusion Tribrid mass spectrometer (ThermoScientific) interfaced with a Dionex UltiMate 3000 Binary RSLCnano System. Peptides were separated onto an analytical C18 column (100μm ID x 25 cm, 5 μm, 18Å Reprosil-Pur C18-AQ beads from Dr Maisch, Ammerbuch-Entringen, Germany) at a flow rate of 350 nl/min. Gradient conditions were: 3%-22% B for 40 min; 22%-35% B for 10min; 35%-90% B for 10 min; 90% B held for 10 min (solvent A, 0.1 % formic acid in water; solvent B, 0.1% formic acid in acetonitrile). The peptides were analyzed using a data-dependent acquisition method, Orbitrap Fusion was operated with measurement of FTMS1 at resolutions 120,000 FWHM, scan range 350-1500 m/z, AGC target 2E5, and maximum injection time of 50 ms; During a maximum 3-second cycle time, the ITMS2 spectra were collected at rapid scan rate mode, with HCD NCE 34, 1.6 m/z isolation window, AGC target 1E4, maximum injection time of 35 ms, and dynamic exclusion was employed for 20 seconds.

The raw data files were processed using ThermoScientific Proteome Discoverer software version 1.4, spectra were searched against the Uniprot-*Mus musculus* (taxonomy id: 10090) or *Schistosoma mansoni* (taxonomy id: 6183) database using Sequest. Trypsin was set as the enzyme with maximum missed cleavages of 2. The precursor ion tolerance was set 10 ppm; MS/MS tolerance 0.8 Da. Carbamidomethylation on cysteine residues was used as static modification; oxidation of methione as well as phosphorylation of serine, threonine and tyrosine was set as variable modifications. Search results were trimmed to 1% FDR for strict and 5% for relaxed condition using Percolator.

### In silico modeling of protein-protein interaction

The amino acid sequence of MEG-8.2 was used to predict its protein structure, with residues 1–20 (N-terminal signal peptide) excluded after identification using PrediSI (http://www.predisi.de/). A 3D model of MEG-8.2 was generated using the Phyre² online server (Protein Homology/analogy Recognition Engine V 2.0) (*77*). The generated model was submitted to the YASARA Energy Minimization Server (*78*) and evaluated using multiple structural validation tools: RAMPAGE (Ramachandran Plot Assessment) (*79*), ProSA-web (Protein Structure Analysis) (*80, 81*), PROCHECK (*82*), ERRAT (*83*), and Verify 3D (*84, 85*). The CD98hc-LAT1 complex was downloaded from the Protein Data Bank (PDB) (*35*) under the accession number 6JMQ. Protein-protein docking analyses were performed using the ClusPro online server (https://cluspro.org), which allows primary usage by submitting two files in PDB format (*86–88*). All models and protein-protein docking visualizations were carried out using PyMOL (v3.0, The PyMOL Molecular Graphics System, Schrödinger, LLC). For the Cct2-2-MEG 8.2 and MEMO-1-MEG 8.2 models, the structures were generated following the same methodology previously described. The predicted structures of CCT-2 and MEMO-1 were obtained from the AlphaFold protein structure database CCT-2 (https://alphafold.com/entry/P80314), MEMO-1 (https://alphafold.com/entry/Q91VH6)]. Similarly, the interaction model for Lactoferrin-MEG 8.2 was generated using the same approach, with the predicted structure of Lactoferrin retrieved from the Protein Data Bank (PDB) under accession number 1LFG (*89*).

### Dot blot assay

Plasma samples were isolated from infected (∼7-weeks post-infection) and uninfected Swiss-Webster mice. Circular pieces of PVDF membranes (0.2 µm) were prepared and placed in a 96-well flat-bottom plate. Membranes were first activated with methanol and subsequently washed with deionized water. 50 µL of uninfected and infected plasma samples were added to five separate wells each. 50 µL of 1X PBS was added in a separate well as a blank control. Plates were incubated overnight at 37°C. Following incubation, membranes were washed three times with TBS-T (Tris-buffered saline with Tween-20), blocked for one hour with a blocking buffer, and incubated overnight at 4°C with varying dilutions of the polyclonal anti-MEG-8.2 antibodies we generated (1:2000, 1:1500, 1:1000, 1:500, and 1:100). Following primary antibody incubation, the membranes were washed three times with TBS-T. An HRP-conjugated anti-rabbit secondary antibody (diluted 1:1000) was added and incubated overnight at 4°C. The membranes were washed three times with TBS-T. Detection was performed using DAB (3,3’-diaminobenzidine) substrate solution kit (34002, ThermoScientific) with an incubation period of 5-15 minutes, allowing colorimetric development. The reaction was stopped once the desired signal intensity was achieved, and relative intensity was quantified using ImageJ software.

### Statistical analysis

GraphPad Prism was used to perform appropriate statistical tests. Specific tests are indicated in the figure legends of each figure.

### Oligonucleotides used in this study

Primers used for cloning, RNAi, in situ hybridization and qPCR are listed in **Table S3**.

## Acknowledgments

*Biomphalaria glabrata* (NMRI) exposed to *Schistosoma mansoni* (NMRI) was provided by the Schistosomiasis Resource Center of the Biomedical Research Institute (Rockville, MD) through NIH-NIAID Contract HHSN272201700014I. The authors thank the University of Wisconsin-Madison Biotechnology Center Gene Expression Center & DNA Sequencing Facility for providing library preparation and next generation sequencing services. This work is supported in part by the Clinical and Translational Proteomics Service Center at the University of Texas Health Science Center. We thank Ogochukwu J. Ezeigwe and Fletcher Metz for their assistance in parts of experiments related to this work in the Lee lab and the Newmark lab, respectively. We thank Dr. Ziyin Li and Dr. Yasuhiro Kurasawa in the Li lab for helping us troubleshoot recombinant protein expression and purification. This work was supported by NIH R01AI175079 (J.L.), Dean’s Startup Fund and Just Missed Grant from McGovern Medical School at UTHealth (J.L.), and Rising STAR Award from The University of Texas System (J.L.). J.L. is the recipient of a Morgridge Postdoctoral Fellowship, and P.A.N. is a Howard Hughes Medical Institute investigator.

---

**Figure S1.**
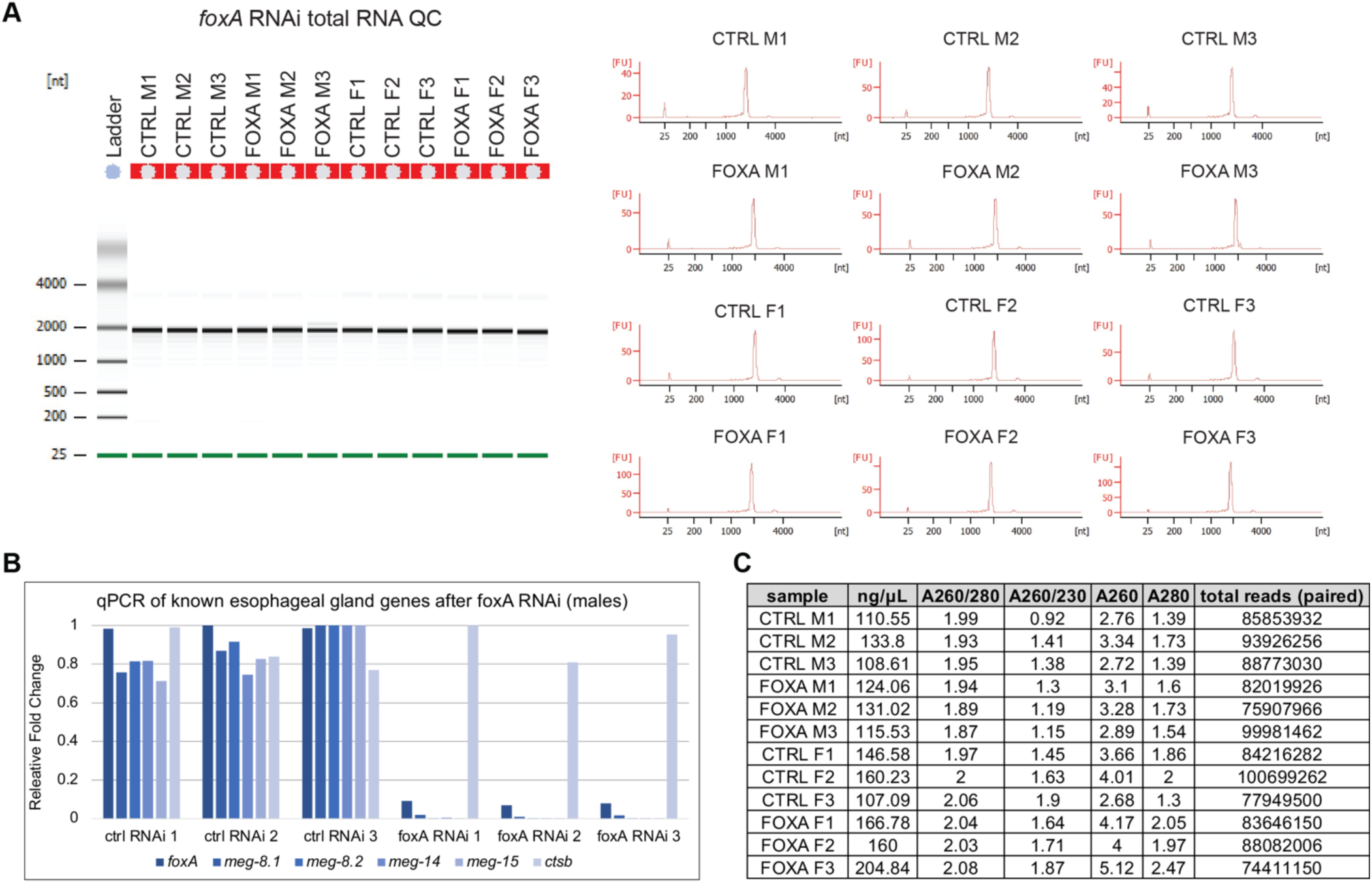
Quality control of total RNA used in RNA-seq. (A) Bioanalyzer results of extracted total RNA samples. (B) qPCR of select known EG genes and non-EG genes (*ctsb*) in cDNA synthesized from the extracted RNA samples. The results show specific downregulation of EG genes in *foxA* knockdown. (C) Summary table of RNA concentration/quality and total reads sequenced for each sample.

**Figure S2.**
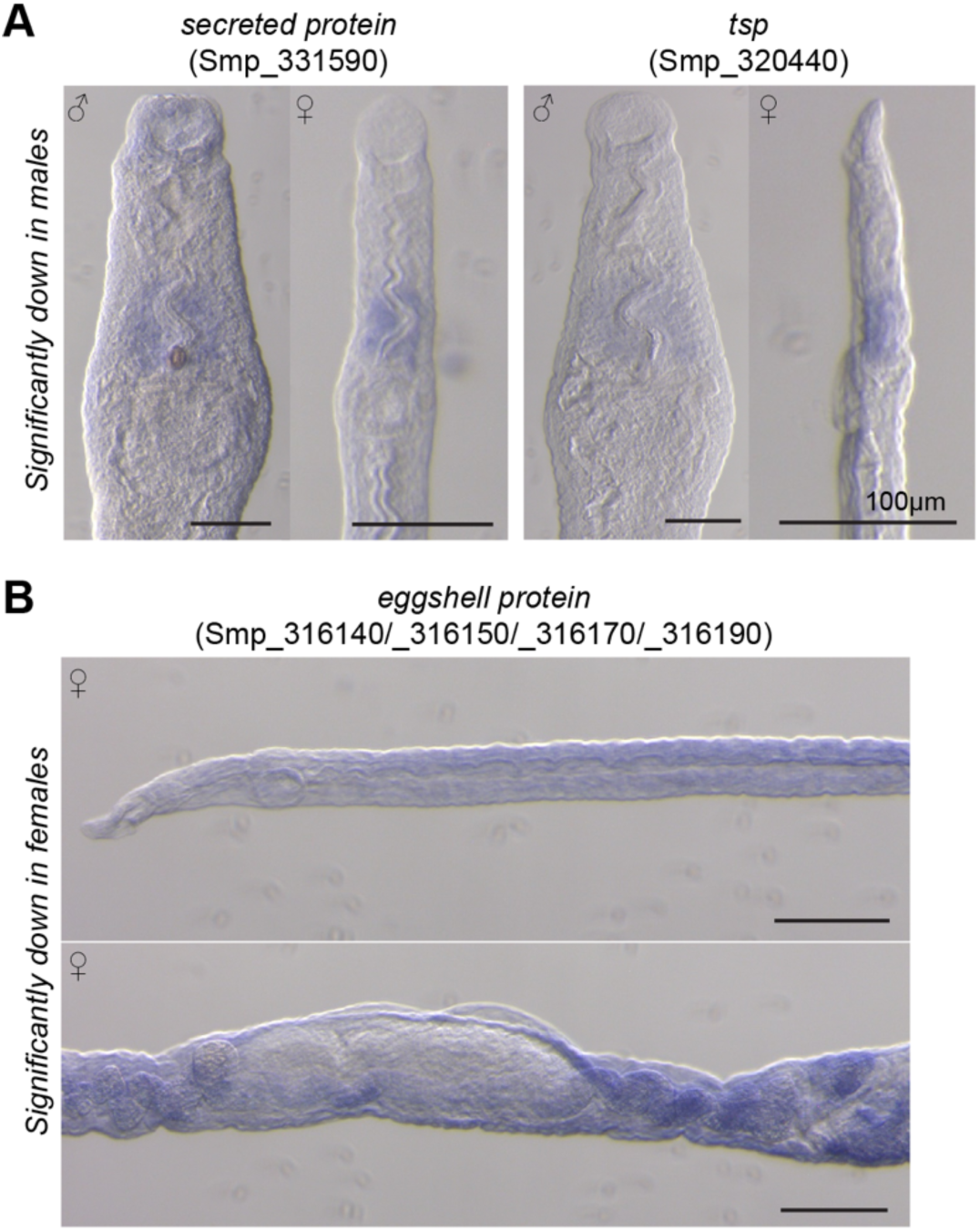
WISH of genes significantly downregulated in males (A) or females (B). (A) Both genes are significantly downregulated only in males but show slight enrichment in both males and females. (B) Eggshell protein downregulated in *foxA* RNAi females is not enriched in the EG but is likely enriched in the accessory reproductive tissues (e.g., vitellaria).

**Figure S3.**
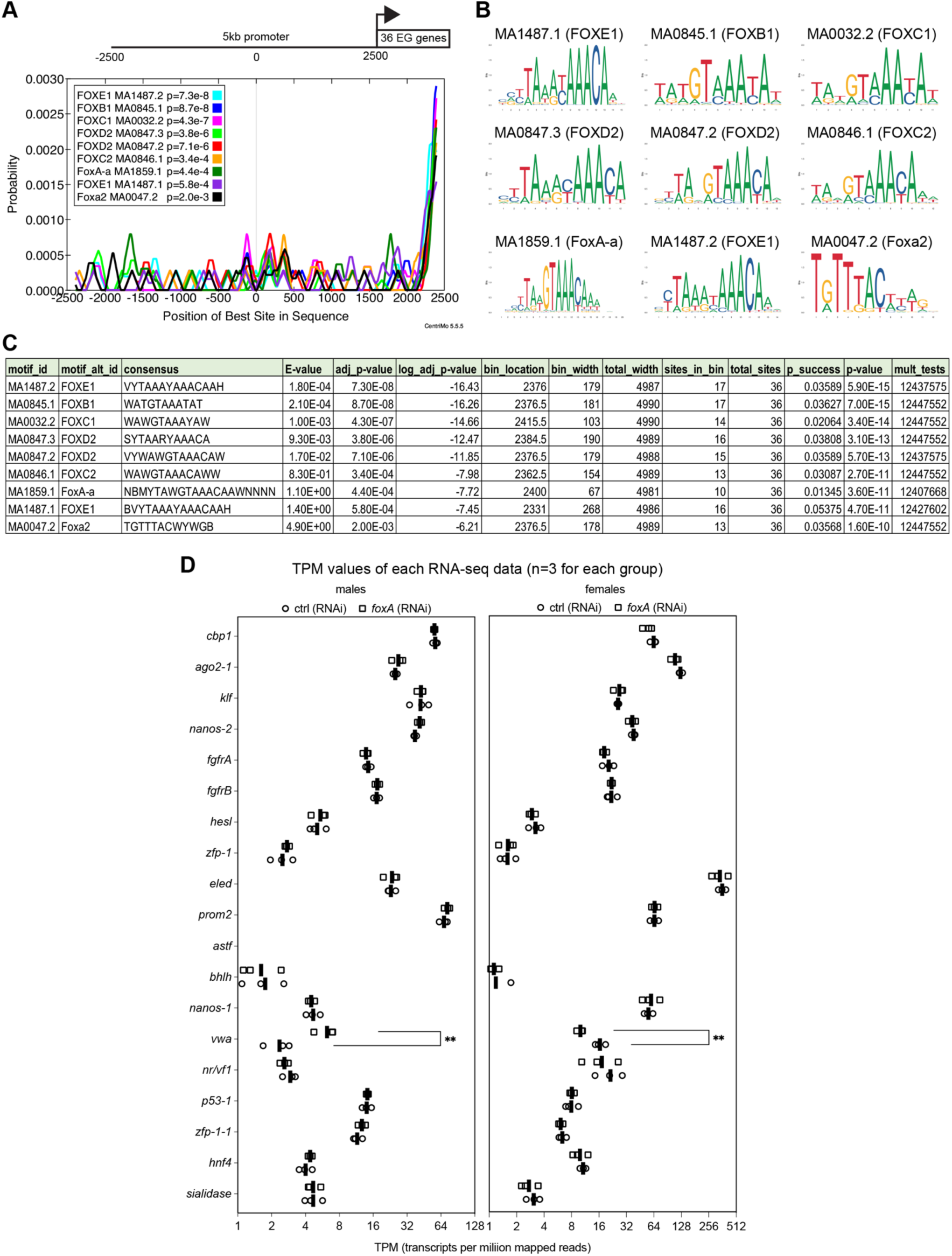
FoxA potentially regulates the EG genes’ transcription directly but has little effect on other cell-type lineages. (A-C) CentriMo analysis of 5kb upstream sequences of 36 EG genes reveals putative forkhead transcription factor binding motifs on most promoters. (A) An overlay of the probability of each motif occurrence. (B) Enriched motifs. (C) A summary table of the location and the significance of each motif. (D) TPM values of cell-type progenitor markers in control and *foxA* RNAi males (left) and females (right). Except for *vwa*, a Mehlis gland marker, expression levels remain largely unchanged.

**Figure S4.**
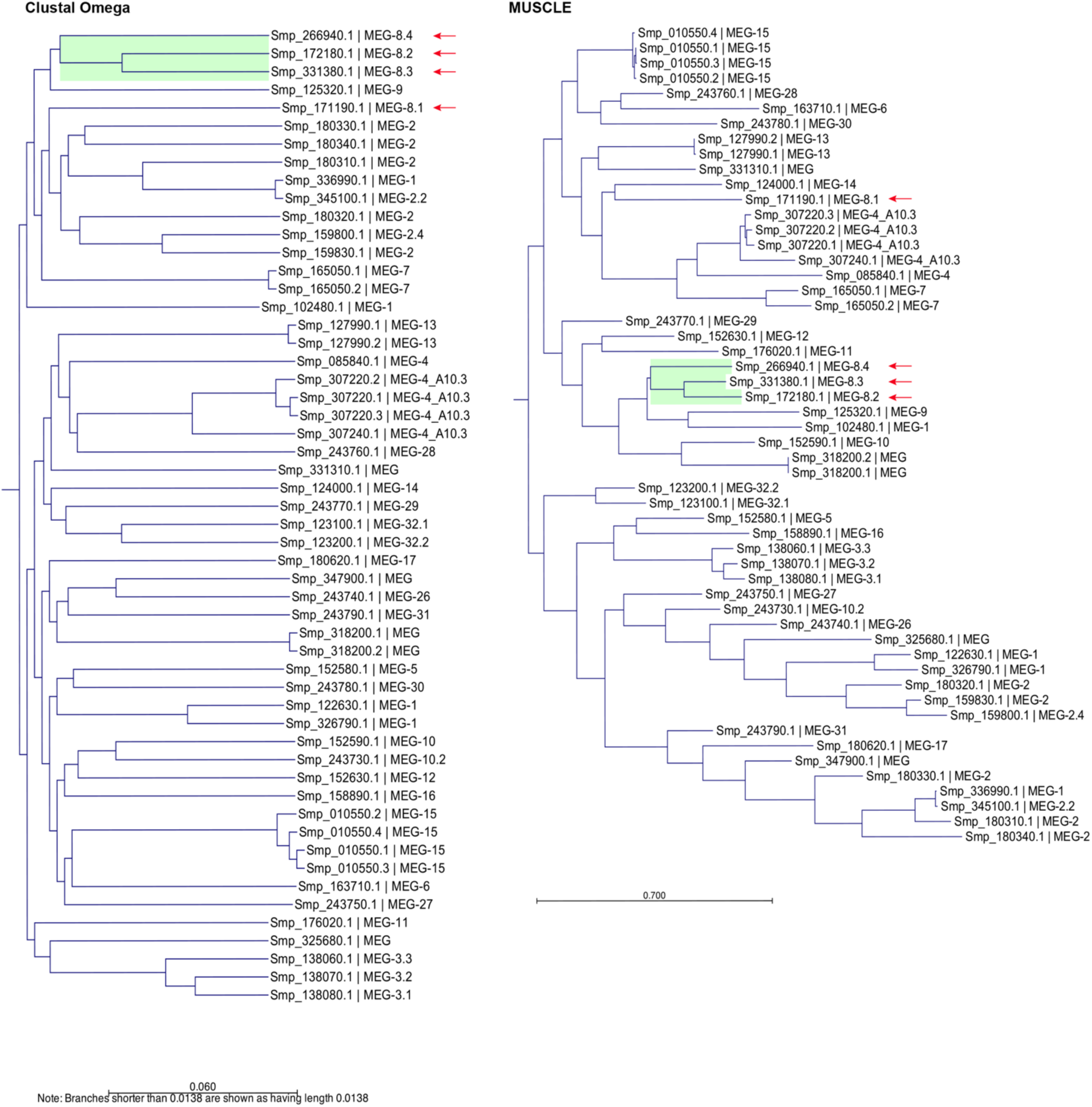
*S. mansoni* MEG family alignment. Left: Clustal Omega (v1.2.0); Right: MUSCLE (Algorithm: Neighbor Joining; Distance measure: Jukes-Cantor; Bootstrap: 100 replicates). Sm-MEG-8 family proteins are indicated with red arrows. Amino acid sequences for all of the proteins were derived from the *S. mansoni* genome (V10) available on WormBase ParaSite.

**Figure S5.**
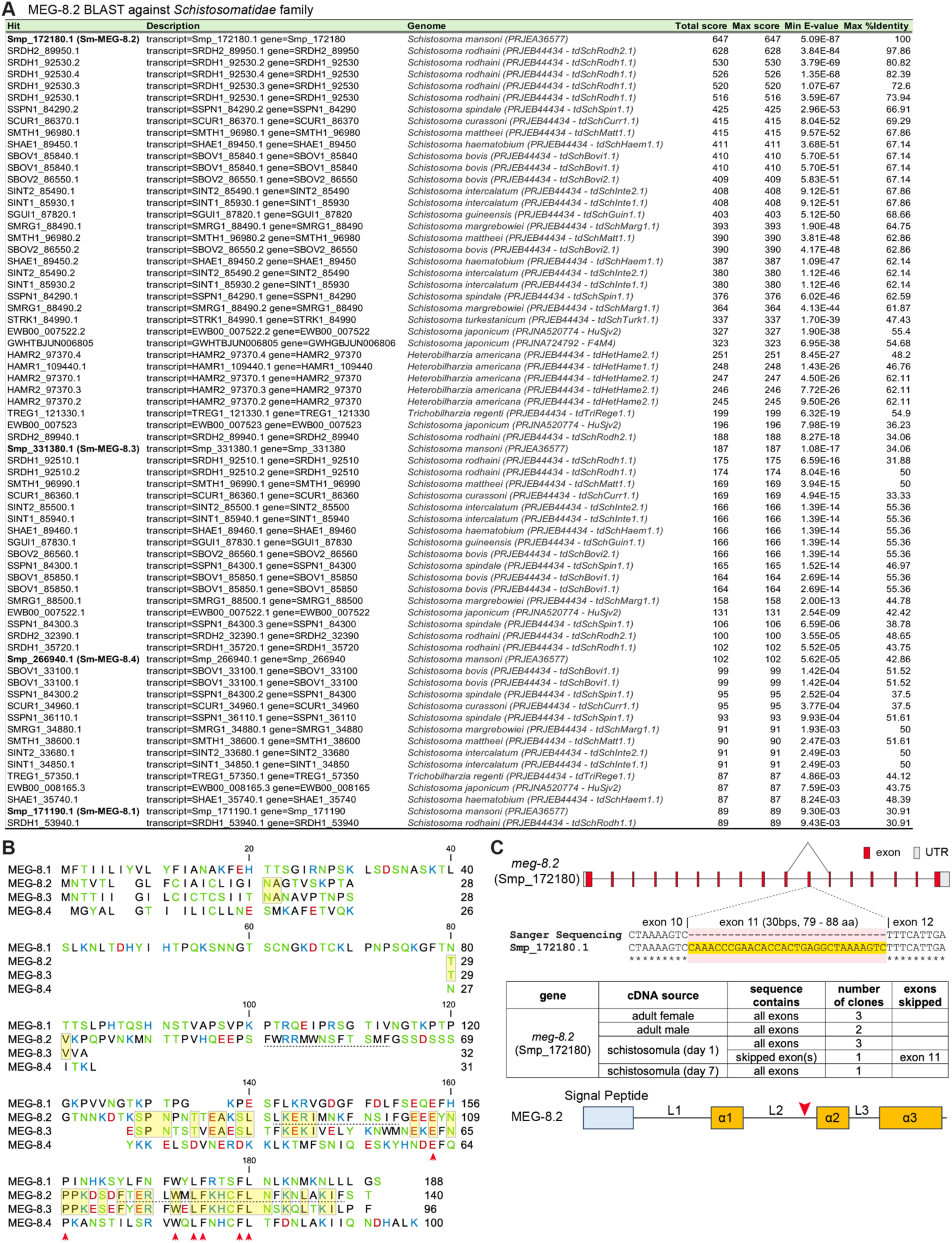
MEG-8 family alignment and a rare case of exon skipping. (A) BLAST of four Sm-MEG-8 proteins across the *Schistosomatidae* family show orthologs in other species. (B) Sm-MEG-8 alignment. The three predicted helices are indicated by a dotted underline. MEG-8.2 and MEG-8.3 share the most residues (yellow box). Residues shared across all four proteins are marked with a red arrowhead. (C) The regional sequence and location of skipped exon 11, which was found in one out of 10 sequenced clones.

**Figure S6.**
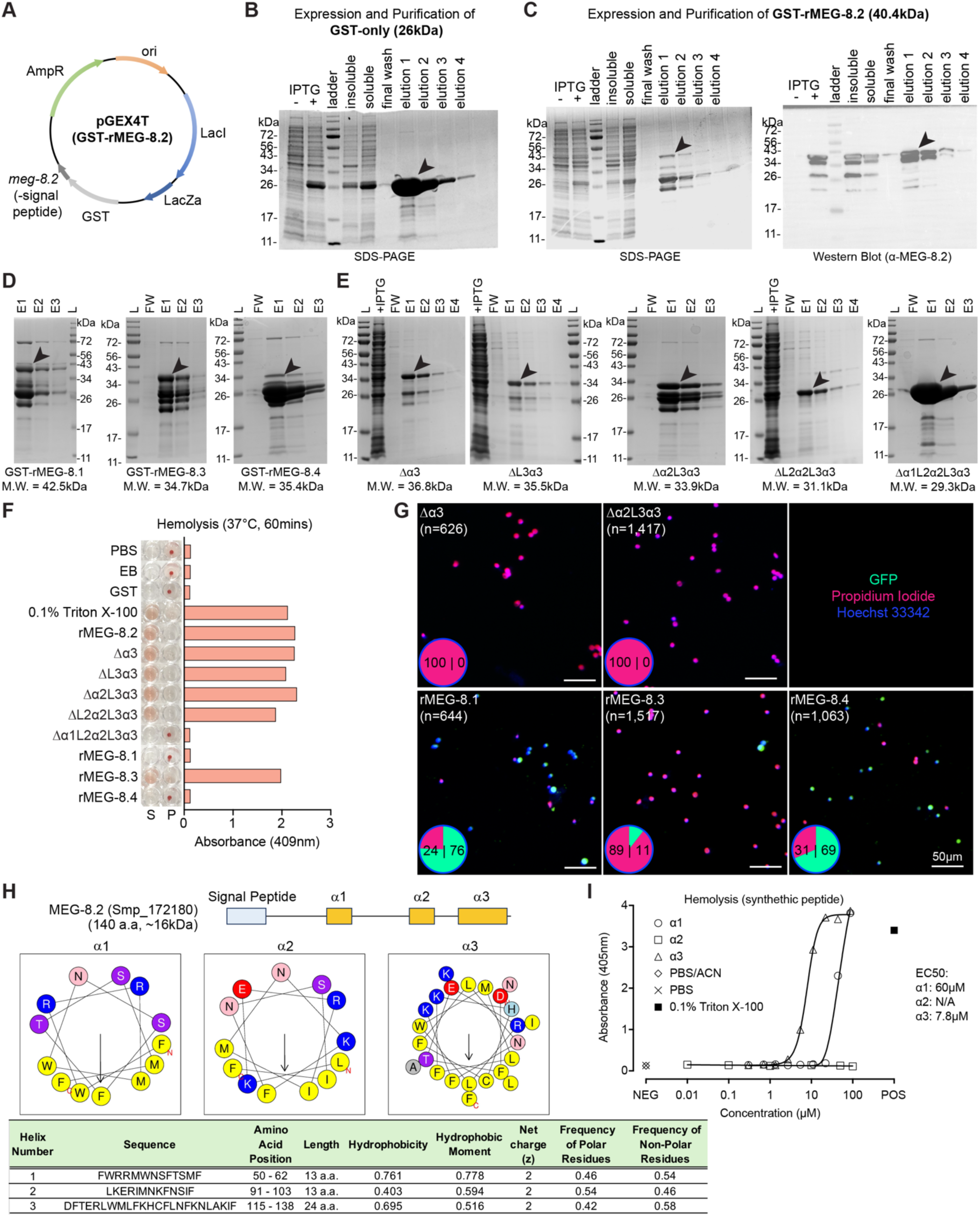
Cell lytic activity of recombinant MEG-8 proteins and mutants. (A) Schematic plasmid map of the bacterial expression vector used to express N-terminal GST-tagged MEG-8 proteins. GST-rMEG-8.2 is shown as an example. (B) SDS-PAGE of GST-only expression and purification steps. (C) Expression and purification of GST-rMEG-8.2. SDS-PAGE (left) shows the highest band corresponding to the expected molecular weight, while a few smaller-size proteins are observed. Western blot (right) using α-MEG-8.2 antibody confirms that these bands are positively labeled, suggesting that while rMEG-8.2 protein is produced, several degradation products are also in the purified mixture. (D) SDS-PAGE of other members of the Sm-MEG-8 family proteins. (E) SDS-PAGE of rMEG-8.2 truncation mutants. (B – D) Arrowheads indicate the expected band size. L: ladder; +IPTG: IPTG induced; FW: final wash; E1 – 4: elution 1 through 4. (F) Hemolysis assay using recombinantly purified MEG-8 proteins. Isolated peripheral blood was treated with indicated proteins for 60 minutes at 37°C. rMEG-8.2 containing α1 region retains the cell lytic activity, as well as rMEG-8.3, but not rMEG-8.1 or rMEG-8.4. EB; elution buffer only. (G) leukocyte lysis by rMEG-8.2 mutants (top) and other MEG-8 family proteins (bottom). GFP-expressing leukocytes were treated with each protein for 10 minutes at 37°C prior to adding PI and Hoechst33342. The pie chart on the lower left corner of each image indicates the viability. n: total number of cells counted. (H) HeliQuest analysis (*90*) of each of the predicted helices shows amphipathic properties, with α1 having the highest hydrophobicity and hydrophobic moment. (I) An independent second experiment of the dose curve of the synthetic peptides. EC50 values are largely in agreement with those shown in Figure 4.

**Figure S7.**
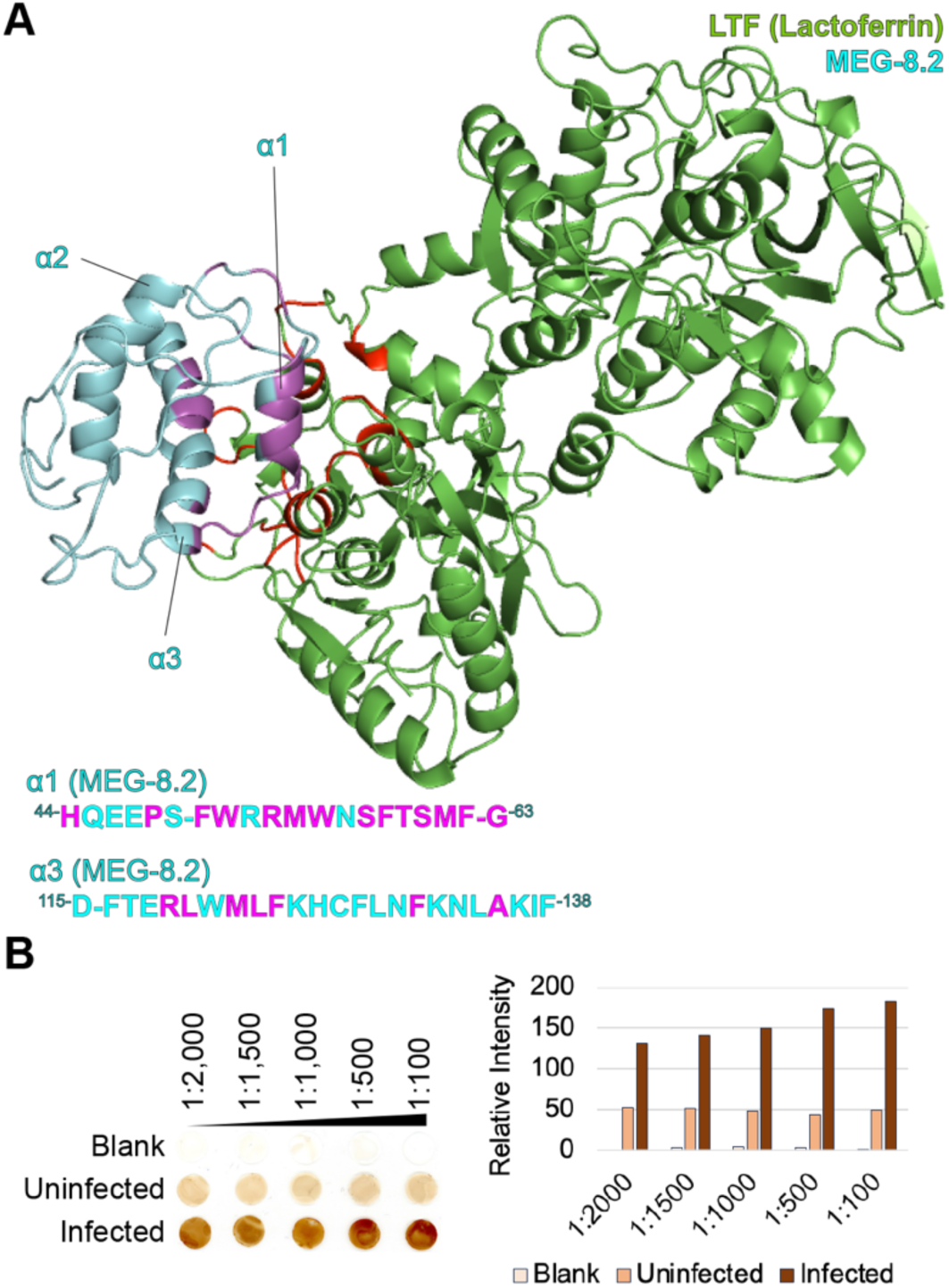
Modeling interaction between LTF and MEG-8.2. (A) MEG-8.2 residues with an interaction distance <4Å are highlighted in magenta. (B) Dot blot of MEG-8.2 in plasma lysate derived from infected and uninfected mice using a range of dilutions of anti-MEG-8.2 antibodies (see Methods).

**Table S1. (separate file)**

Differential expression analysis of *foxA* RNAi RNA-seq.

**Table S2. (separate file)**

LC-MS/MS analysis of rMEG-8.2 pull-down hits.

**Table S3. (separate file)**

List of oligonucleotides used in this study.

**Table S4. (separate file)**

Plasmid sequences for all constructs used for recombinant protein expression.

